# Dimerization of the *S. cerevisiae* Spo11 core complex

**DOI:** 10.64898/2026.01.16.699991

**Authors:** Hajar Aït Bella, Mahesh Survi, Julian Urdiain-Arraiza, Dayishaa Daga, Vijayalakshmi V. Subramanian, Andreas Hochwagen, Corentin Claeys Bouuaert

**Affiliations:** Louvain Institute of Biomolecular Science and Technology, Université catholique de Louvain, 1348 Louvain-La-Neuve, Belgium; Department of Biology, New York University, New York, NY 10003, USA; Department of Biology, IISER Tirupati, Tirupati, 517507, Andhra Pradesh, India

## Abstract

Spo11 initiates meiotic recombination by introducing programmed DNA double-strand breaks. DNA cleavage occurs via a topoisomerase-like mechanism involving hybrid active sites formed at the dimer interface. However, in contrast to its topoisomerase relative (Topo VI), Spo11 does not form a stable dimer, likely to prevent uncontrolled DNA cleavage. Here, we investigated the dimerization of *S. cerevisiae* Spo11 in complex with its partners Ski8, Rec102, and Rec104. We show that the Spo11 complex dimerizes transiently on DNA, forming unstable dimeric complexes with duplex and branched DNA substrates. Guided by AlphaFold modeling of a pre-cleavage complex, we identified mutations that reduce dimerization. Surprisingly, DSB formation is resilient to mutagenesis of the Spo11 dimer interface, implying that additional factors promote dimerization *in vivo*. Finally, we found that Rec102 exerts a key DNA-binding function, essential for catalysis, and show that it also participates in dimerization through *trans* contacts with Ski8. Our work provides new insights into the mechanism of Spo11 dimerization and the role of its partners in initiating meiotic recombination.

## Introduction

Meiotic recombination ensures accurate segregation of homologous chromosomes during gametogenesis and reshuffles allelic combinations between generations to drive genetic diversity and evolution (Hunter 2015; Zickler and Kleckner 2023). Recombination is initiated by Spo11, a topoisomerase-like protein that introduces programmed DNA double-strand breaks (DSBs) throughout the genome (Yadav and Claeys Bouuaert 2021). Although DSB formation by Spo11 is tightly controlled, the mechanisms that govern its catalytic activity remain to be fully elucidated.

Spo11 derives from an ancestral type IIB topoisomerase, Topo VI, which operates as a heterotetramer containing two DNA-cleavage (A) subunits and two ATPase (B) subunits (Corbett et al. 2007; Harper et al. 2025) (**Figure 1A**). The A subunits, homologous to Spo11, cleave DNA via a transesterification mechanism involving a nucleophilic attack from an active site tyrosine to the DNA backbone, thereby producing 5′-phosphotyrosyl covalent intermediates. Two strands are cleaved in concert, producing breaks with 2-nucleotide (nt) 5′-overhangs. Each strand is cleaved by a hybrid active site constituted at the interface between two subunits, one contributing the catalytic tyrosine, the other contributing acidic residues that coordinate divalent metal ions essential for catalysis. Recent *in vitro* reconstitution experiments demonstrated that Spo11 shares these mechanistic features with Topo VI (Oger and Claeys Bouuaert 2025b; Oger and Claeys Bouuaert 2025a; Tang et al. 2025; Zheng et al. 2025), as had previously been inferred from *in vivo* work (Keeney 2001). Hence, just like Topo VI, DNA cleavage by Spo11 necessarily requires dimerization.

**Figure 1:**
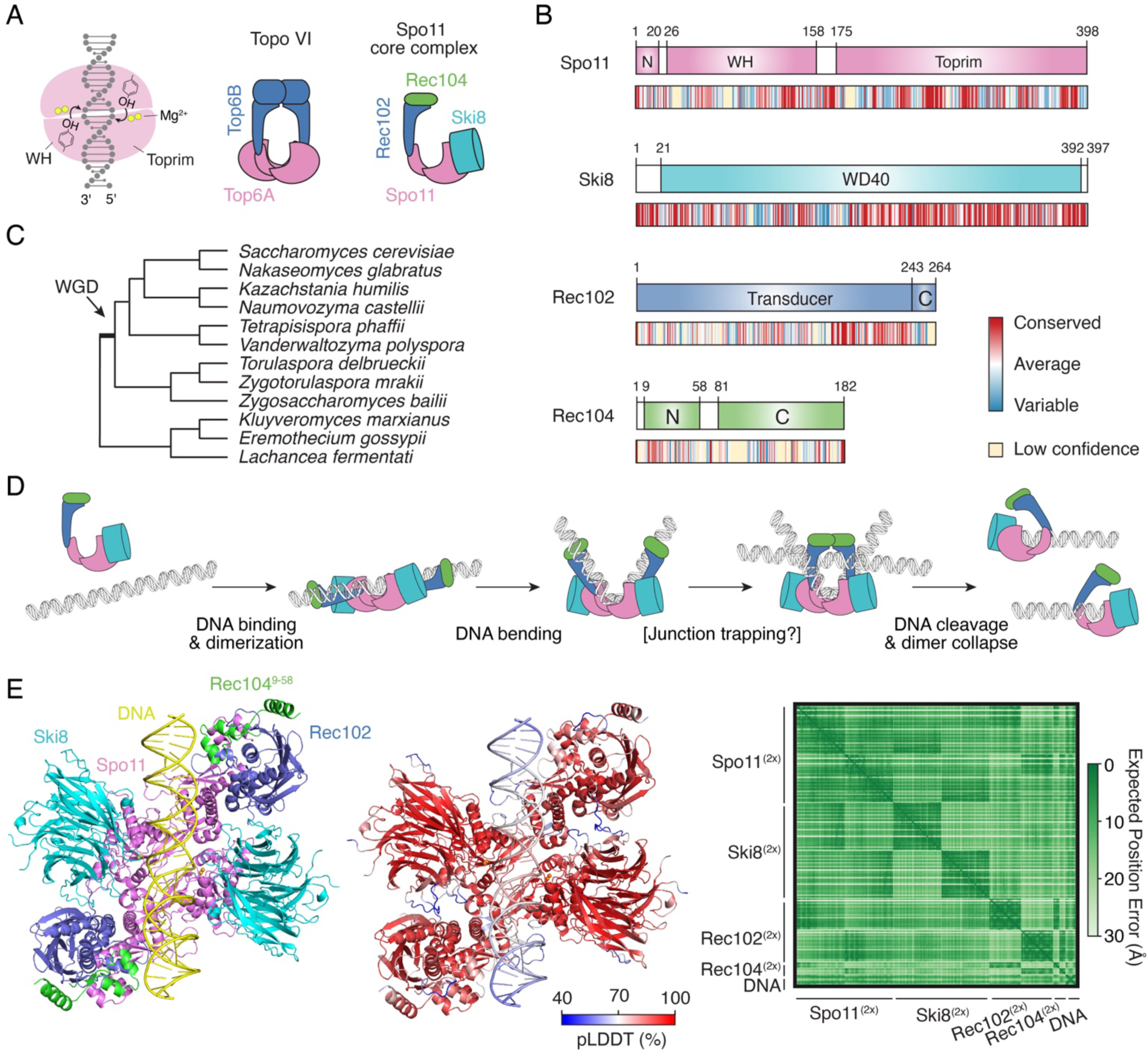
The *S. cerevisiae* Spo11 core complex. **(A)** Chemistry of strand cleavage and comparison of Topo VI and the Spo11 core complex. **(B)** Domain structure of core complex proteins Spo11, Ski8, Rec102, and Rec104. Sequence conservation across Saccharomycetaceae is shown (see **Supplementary Figures 3-6**). **(C)** Phylogenetic tree of Saccharomycetaceae species used for sequence conservation analyses. Topology based on (Wolfe et al. 2015; Liu et al. 2024). WGD, whole-genome duplication. **(D)** Cartoon of the proposed dynamics of Spo11 association with DNA (Claeys Bouuaert et al. 2021b). The core complex is monomeric (1:1:1:1 stoichiometry) in solution. The complex may bind DNA as a monomer and dimerize on DNA, upon which Spo11 induces a sharp DNA bend. The complex may also trap a second DNA duplex prior to DNA cleavage. Catalysis is followed by a collapse of the dimer interface, producing irreversible breaks. (**E**) Structural modeling of a dimeric Spo11 core complex (2:2:2:2 Spo11:Ski8:Rec102:Rec104^9-58^) bound to DNA and four Mg^2+^ ions. Rec104 was trimmed (residues 9-58) to match the resolved region in the monomeric Spo11 core complex cryo-EM structure (PDB: 8URQ (Yu et al. 2024)). Left, AlphaFold3 model colored by chain. Spo11 (violet), Ski8 (cyan), Rec102 (blue), Rec104 (green), DNA (yellow). Center, model colored by confidence score (ipTM 0.85, pTM 0.86, global pLDDT 82). Right, predicted alignment error plot, showing high confidence in the structure and relative position of all subunits.

In striking contrast with Topo VI, however, purified Spo11 is monomeric in solution. This was shown for the *S. cerevisiae* (Claeys Bouuaert et al. 2021b), *M. musculus* (Oger and Claeys Bouuaert 2025b; Tang et al. 2025; Zheng et al. 2025), and *C. elegans* (Yeh et al. 2017) orthologs, suggesting that it is a conserved feature of Spo11 that could play a crucial role in controlling catalysis. *M. musculus* and *C. elegans* SPO11/SPO-11 have been purified by themselves, without any additional partners. While purified mouse SPO11 was active *in vitro* (Oger and Claeys Bouuaert 2025b), the worm ortholog was not (Yeh et al. 2017). Yeast Spo11 required partners Rec102, Rec104 and Ski8 for purification, together forming a 1:1:1:1 complex (Claeys Bouuaert et al. 2021b). This complex, referred to as the core complex, is also catalytically inactive *in vitro*.

The B-type subunit derived from the ancestral topoisomerase is a conserved and essential member of the meiotic DSB machinery in most eukaryotes, although the ATP binding and hydrolysis activities are not conserved (Robert et al. 2016; Vrielynck et al. 2016). In yeast, the B-type subunit, Rec102, entirely lacks the ATPase GHKL domain (Robert et al. 2016) (**Figure 1A-C**). This domain was replaced by Rec104, a poorly conserved and poorly structured subunit whose function remains enigmatic (Claeys Bouuaert et al. 2021b; Yu et al. 2024). The last member of the *S. cerevisiae* Spo11 core complex, Ski8, also has a separate function as part of the Ski complex that channels mRNA to the exosome for degradation (Arora et al. 2004; Halbach et al. 2013). In many fungi, Ski8 evolved as a *bona fide* Spo11 partner, although the mechanism whereby it supports Spo11 activity remains unclear (Tesse et al. 2003; Steiner et al. 2010).

Even though the yeast Spo11 core complex is catalytically inactive *in vitro*, it binds DNA efficiently. Atomic force microscopy (AFM) analyses showed that the complex binds internally to DNA duplexes, its presumed substrate, and appears to induce sharp DNA bends (Claeys Bouuaert et al. 2021b). The core complex also frequently locates at sites where two DNA duplexes cross each other. This activity is shared with type II topoisomerases, which is presumably relevant to their function in modulating DNA topology (Alonso-Sarduy et al. 2011; Timsit 2011; Wendorff and Berger 2018). Double-strand break formation by Spo11, on the other hand, does not require strand passage, and the relevance of this junction-binding activity remains unclear. Finally, the Spo11 core complex exhibits a particularly tight binding activity to double-stranded DNA ends with 2-nt 5′-overhangs that mimic the product of DNA cleavage (Claeys Bouuaert et al. 2021b).

We previously proposed that these DNA-binding activities reflect intermediates in a multi-step reaction pathway (Claeys Bouuaert et al. 2021b), inspired by and adapted from the mechanism of Topo VI (Corbett et al. 2007; Wendorff and Berger 2018). The hypothesis is that monomeric (1:1:1:1) core complexes bind duplex DNA substrates and, at some frequency, may dimerize on DNA (**Figure 1D**). Dimerization would be accompanied by the induction of a ∼100-120° DNA bend, upon which the complex would perhaps trap a second DNA duplex. After DNA cleavage, the dimer interface of Spo11 would collapse and the complex would relax on the DNA ends in a stable post-cleavage state that prevents reaction reversal (Claeys Bouuaert et al. 2021b; Oger and Claeys Bouuaert 2025a).

A recent cryo-EM structure of the *S. cerevisiae* Spo11 core complex bound to a double-stranded DNA end provided detailed structural insights into the presumed post-cleavage state (Yu et al. 2024). On the other hand, no structural information on pre-cleavage complexes is currently available. Hence, the conformational dynamics and molecular interactions involved during catalysis remain to be fully elucidated.

Here, we performed a structure-function analysis of the *S. cerevisiae* Spo11 core complex to delineate the role of the Spo11 dimer interface and the molecular functions of Spo11 partners. We find that the core complex dimerizes to form a transient, unstable, pre-cleavage complex.

Mutations that compromise Spo11 dimerization *in vitro* lead to remarkably subtle phenotypes *in vivo*, highlighting the resilience of the meiotic DSB machinery. Finally, we show that Rec102 directly contacts DNA prior to DSB formation and participates in dimerization through *trans* contacts with Ski8. This work provides new insights into the molecular mechanism of meiotic DSB formation in *S. cerevisiae*.

## Results

### Modeling of a pre-cleavage Spo11 core complex

To investigate the assembly of pre-cleavage complexes, we modeled the structure of a dimeric Spo11 core complex bound to DNA using AlphaFold3 (Abramson et al. 2024). As we previously showed for *Mus musculus* SPO11 (Oger and Claeys Bouuaert 2025b), AlphaFold3 fails to predict the structure of a dimeric Spo11 complex in the absence of DNA (**Supplementary Figure 1A**). In the presence of DNA, the confidence of the relative positioning of two Spo11 complexes was significantly improved (**Supplementary Figure 1B**). The sequence of the DNA substrate used for modeling also impacted the model quality, although sequences that match a Spo11 consensus cleavage site were not optimal (**Supplementary Table 1**).

The structures of Spo11, Ski8 and Rec102 were predicted with high confidence and were generally in line with the published cryo-EM structure of the core complex in a post-cleavage state (Yu et al. 2024) (**Supplementary Figure 2**), and the interaction surfaces between subunits were accurately recapitulated (see comparisons of predicted and experimental interface residues on sequence alignments in **Supplementary Figures 3-6**). On the other hand, the structure of Rec104 was not predicted with confidence (**Supplementary Figure 1**), consistent with the fact that only the N-terminal ∼60 residues of Rec104 are resolved in the cryo-EM maps, likely due to structural heterogeneity of the ∼120 amino-acid C-terminal part (Yu et al. 2024).

Optimizing the DNA sequence and truncating Rec104 (residues 9 to 58) yielded a pre-cleavage model with high confidence scores (ipTM 0.85, pTM 0.86, global pLDDT 82) (**Figure 1E**). In the model, the DNA substrate is predicted to be bent at an angle of 100°, consistent with previous AFM analyses (Claeys Bouuaert et al. 2021b) and similar to AlphaFold predictions of Spo11 complexes from other species (Oger and Claeys Bouuaert 2025b). The catalytic tyrosines are located about 6.5 Å away from the scissile phosphates and 8 Å from the catalytic metal ions across the dimer interface. Hence, this model likely provides a relatively accurate representation of a pre-cleavage Spo11 core complex, although the active sites are not poised for catalysis.

### The Spo11 core complex dimerizes transiently on DNA

To understand how Spo11 engages with its DNA substrate, we sought to study two poorly characterized binding modes of the Spo11 core complex: internal binding to DNA duplexes and binding of DNA crossings (junctions). To this end, we analyzed the DNA-binding activity of purified core complexes by gel shift assay using radiolabeled substrates assembled with 40-nt complementary oligonucleotides. The substrates contained either a 40-bp DNA duplex (dsDNA) or a four-way branched structure with 20-bp arms (Holliday Junction, HJ), which mimics configurations where two DNA duplexes cross each other (Wendorff and Berger 2018). To minimize end binding, the substrates harbored 1-nt 3′-extensions that strongly reduce the end-binding activity of the Spo11 core complex (Claeys Bouuaert et al. 2021b).

We titrated the protein in the presence of the labeled DNA substrates and separated complexes by native gel electrophoresis (**Figure 2A**). The Spo11 complex bound the dsDNA and HJ substrates with an apparent affinity of 1.2 nM and 1.3 nM, respectively (**Figure 2B**), which corresponds to about a 20-fold lower affinity than that for double-stranded DNA ends with 2-nt 5′-overhangs (K_D_ ≈ 0.06 nM) (Claeys Bouuaert et al. 2021b).

**Figure 2:**
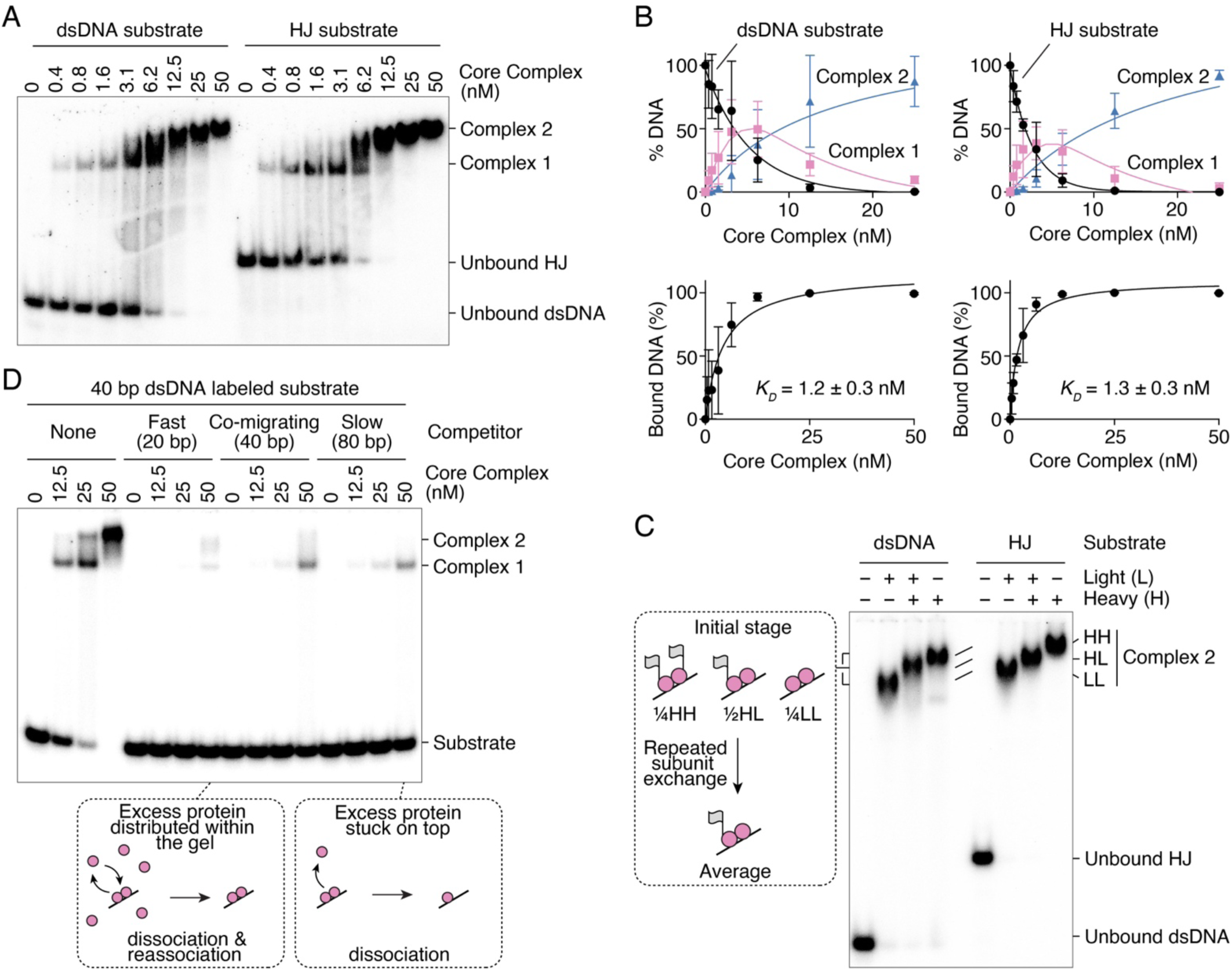
The Spo11 core complex dimerizes on DNA to form an unstable complex. **(A)** Gel shift analysis of the binding activity of the core complex to 40 bp duplex DNA (dsDNA) and 4-way branched DNA with 20 bp arms (Holliday Junction, HJ). Both substrates have DNA ends with 1-nt 3′-overhangs to minimize end-binding (Claeys Bouuaert et al. 2021b). The migration of Complex 2 decreases at higher protein concentration, presumably due to faster re-association of complexes that dissociate during electrophoresis, or due to weak non-specific interactions with additional core complexes. **(B)** Quantification of gel shift analyses of core complex binding to dsDNA and HJ substrates (mean ± SD of n = 3 independent experiments). **(C)** Gel shift analysis with combinations of core complexes (100 nM total) with (heavy) and without (light) MBP-YFP (grey flags) fused to Rec102. **(D)** Gel shift analysis of DNA complexes with a radiolabeled 40 bp linear dsDNA substrate (1 nM) in the presence of 25-fold excess (in base pairs) of fast-migrating (20 bp), co-migrating (40 bp), or slow-migrating (80 bp) competitor.

Binding of the core complex to the dsDNA and HJ substrates produced two complexes, referred to as Complex 1 and Complex 2 (**Figure 2A**). Complex 1, of higher electrophoretic mobility, presumably contains a single heterotetrameric Spo11 core complex, while the slower-migrating Complex 2 likely contains two core complexes. Complex 1 peaked at about 3-6 nM protein, with ∼25% unbound substrate and ∼25% in Complex 2 (**Figure 2B**). This is close to the expected distribution if both complexes bind the substrate independently. Hence, this provides little evidence that the core complex cooperatively dimerizes on DNA.

To verify the stoichiometry of Complex 1 and Complex 2, we analyzed DNA binding with purified core complexes that either had an MBP and a YFP tag on the N-terminus of Rec102 (heavy) or did not (light). Both protein complexes produced similar binding patterns, except that the complexes assembled with light proteins had higher electrophoretic mobility than the heavy ones. If Complex 2 contains two copies of the core complex, mixing the heavy and light versions should produce three bands: heavy-heavy, light-light and heavy-light dimers, expected at a 1:1:2 ratio. Samples that contained a mixture of heavy and light proteins indeed produced a band of intermediate mobility between the heavy and light Complex 2 (**Figure 2C**). However, in that sample, the bands corresponding to heavy-heavy and light-light Complex 2 were absent from the gel. This could either indicate that the heavy and light proteins preferentially heterodimerize, or that the complexes undergo significant subunit exchange during electrophoresis such that, regardless of the initial distribution at the time of loading the gel, the complexes repeatedly redistribute within the gel and yield, on average, one heavy copy and one light copy.

To investigate this potential dynamic reassociation during electrophoresis, we performed a gel shift assay in the presence of competitor substrates of different length that either migrated ahead of (20 bp), co-migrated with (40 bp), or migrated behind (80 bp) the radioactive dsDNA substrate. The rationale was that the short competitor would produce fast-migrating invisible complexes that would dissociate during electrophoresis, thereby leaving unbound proteins ahead of the radiolabeled complex. This would provide an opportunity for subunit exchange, facilitating reassembly of Complex 2 in case it collapses during electrophoresis. In contrast, the longer competitor DNA would produce slow-migrating invisible complexes and trap proteins behind the radiolabeled complex. In that case, a labeled Complex 2 that collapses during electrophoresis would encounter few unbound proteins, preventing reassembly. As predicted, Complex 2 was abundant in the presence of the fast-migrating competitor, but almost undetectable with the slower-migrating competitor (**Figure 2D**). We conclude that Complex 2 contains two copies of the core complex, but that it is unstable and experiences rapid subunit exchange during electrophoresis.

### Mutagenesis of the Spo11 dimer interface

To further investigate Spo11 dimerization, we analyzed the dimer interface predicted by AlphaFold and identified residues potentially involved in dimerization, including K126, N127, S136, V138, E139, F203, K206, P207, T239, K240, N243 (**Figure 3B**). To search for a dimerization-defective mutant, we mutated these residues alone or in combination and established their effect on meiotic recombination using a heteroallele recombination assay. In this assay, *spo11Δ* diploid strains that harbor two defective copies of *arg4* with inactivating mutations at opposite extremities are transformed with complementation vectors expressing wild-type or mutant Spo11. Following a few hours of meiotic induction by nitrogen depletion to trigger homologous recombination, strains are plated on medium lacking arginine and reverted to vegetative growth. Recombination between the defective *arg4* alleles can restore a wild-type *ARG4* gene, allowing cells to grow in the absence of arginine. The frequency of *ARG+* colonies therefore provides a semi-quantitative measure of recombination, and therefore DSB formation. To our surprise, among nine mutant alleles tested, none yielded a significant recombination defect (**Figure 3C**).

**Figure 3:**
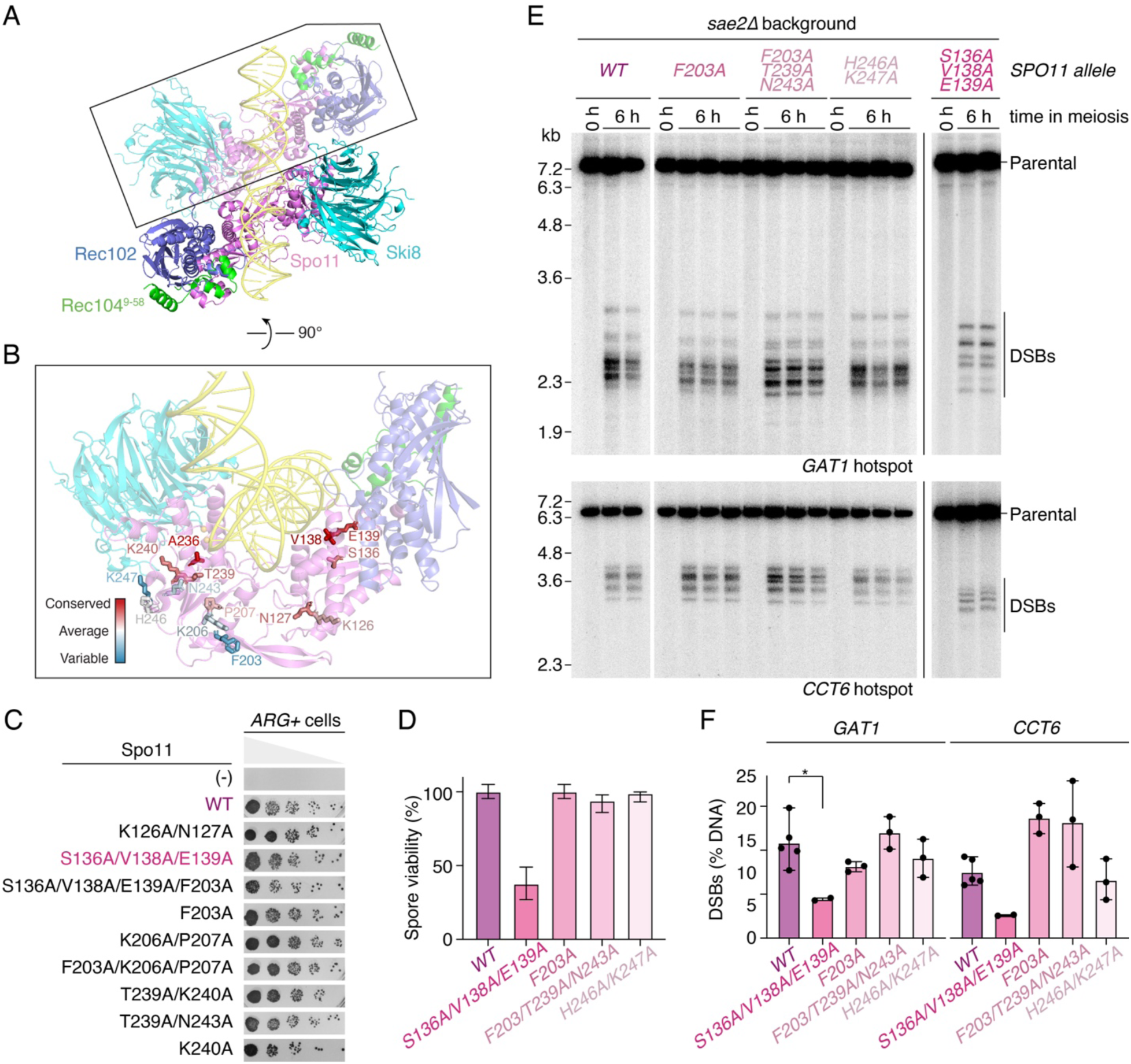
The Spo11 dimer interface is highly tolerant to mutagenesis. **(A)** AlphaFold model of a 2:2:2:2 core complex bound to a 40 bp dsDNA substrate. Spo11 (violet), Ski8 (cyan), Rec102 (blue), Rec104 (residues 9-58, green). **(B)** Spo11 residues at the predicted dimer interface, colored based on their conservation score among Saccharomycetaceae. **(C)** Mutagenesis analysis of predicted Spo11 dimerization mutants by *arg4* heteroallele recombination assay. Five-fold serial dilutions are shown. **(D)** Spore viability of predicted Spo11 dimerization mutants. n = 80 spores per strain. Error bars at 95% confidence intervals of the proportion. **(E)** Southern blot analysis of DSB formation at the *GAT1* and *CCT6* hotspots in wild-type and *spo11* mutant strains in a *sae211* background. For each strain, two or three biological replicates are shown. **(F)** Quantification of gels in panel E (mean ± range of 2-5 biological replicates per strain).

Since the *arg4* heteroallele recombination assay does not provide a sensitive measure of DSB formation, we selected four alleles (*S136A/V138A/E139A*, *F203A*, *F203A/T239A/N243A*, *H246A/K247A*) and introduced these mutations at the endogenous *SPO11* locus by gene replacement. Among those, only the *spo11-S136A/V138A/E139A* (SVE) mutant showed reduced spore viability (37.5%, **Figure 3D**). To characterize the effect of the mutations on DSB formation, we performed Southern blot analysis at the *GAT1* and *CCT6* hotspots in a *sae2Δ* background that accumulates unresected breaks. As compared to the wild type, all the mutants showed similar DSB levels and distribution, except for the *spo11-SVE* mutant that showed reduced and redistributed breaks, in particular at the *GAT1* hotspot (**Figure 3E, F**). The impact of the mutations on the position of cleavage sites likely reflects a reduced efficiency in the assembly of a pre-cleavage complex, as was previously observed for a *spo11-F260A* mutant that has reduced DNA-binding activity (Diaz et al. 2002; Claeys Bouuaert et al. 2021b).

To verify the impact of the Spo11-SVE mutation on complex assembly, we purified the mutant core complex and analyzed its DNA-binding activity on linear and branched substrates with 1-nt 3′-overhangs to assess duplex- and junction-binding activities, and on a short hairpin substrate with a 2-nt 5′-overhang to assess its end-binding activity. The Spo11-SVE mutation did not significantly affect the affinity of the core complex for any of the DNA substrates (**Figure 4A, B**). However, the formation of Complex 2 was strongly reduced, consistent with reduced dimerization (**Figure 4C**).

**Figure 4:**
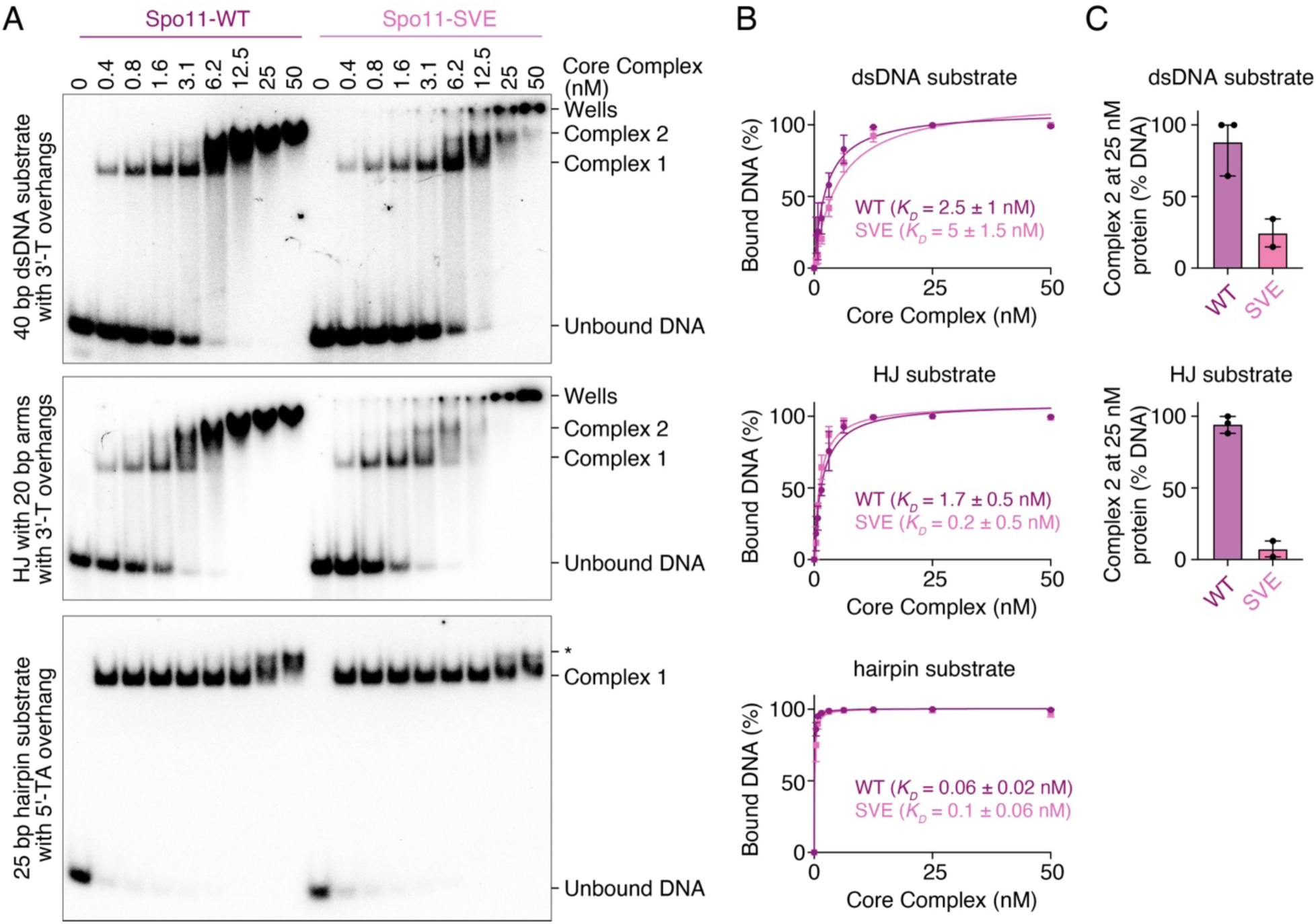
The Spo11-SVE mutation compromises core complex dimerization on DNA. **(A)** Gel shift analysis of the binding activity of wild-type core complexes and Spo11-S136A/V138A/E139A (SVE) mutants to DNA duplexes (40 bp dsDNA with 1-nt 3′-overhangs), Holliday junctions (HJ with 20 bp arms and 1-nt 3′-overhangs), or DNA ends (25 bp hairpin substrates with a 2-nt 5′-overhang). The band labeled * corresponds to monomeric complexes likely bound to opposite ends of the hairpin DNA substrate. The Spo11-SVE mutant produces more signal in the wells than the wild type, presumably because the mutations cause increased exposure of hydrophobic surfaces that lead to aggregation. **(B)** Quantification of the affinity of wild-type and SVE mutants for each substrate. **(C)** Quantification of Complex 2 at 25 nM core complexes (mean ± range of three (WT) or two (SVE) independent experiments).

We also purified core complexes harboring the Spo11-F203A/T239A/N243A (FTN) and Spo11-H246A/K247A (HK) mutations. Although these mutations do not cause a detectable phenotype *in vivo*, both lead to reduced Complex 2 assembly *in vitro*, confirming that the mutations indeed target the dimer interface (**Supplementary Figure 7**). Since Spo11-FTN, HK, and SVE mutants have different *in vivo* phenotypes despite similar *in vitro* defects, this suggests that parts of the dimer interface are readily stabilized by external factors, while others are less so.

We conclude that the dimer interface of Spo11 is highly resilient to mutagenesis, suggesting that additional factors stabilize Spo11 dimers *in vivo*. Nevertheless, Spo11 residues S136, V138 and E139 are important for dimerization *in vitro* and DSB formation *in vivo*.

### Conservation of the Spo11 dimer interface

Given the key role of Spo11 dimerization in catalysis (Oger and Claeys Bouuaert 2025b; Zheng et al. 2025), the tolerance of the dimer interface to mutagenesis is somewhat counter-intuitive. To better understand this, we performed an evolutionary analysis to evaluate the conservation of the Spo11 dimer interface compared to that of Top6A.

We sought to test the hypothesis that the Spo11 dimer interface would have experienced relaxed selective pressure compared to that of Top6A. This would account for the high mutation tolerance we observed, and would also explain why AlphaFold consistently fails to predict the structure of dimeric Spo11 complexes in the absence of DNA, while that of Top6A is predicted with confidence (**Supplementary Figure 8A**). Indeed, relaxed selective pressure may preclude the co-evolutionary analysis that AlphaFold relies upon to constrain the structural model (Abramson et al. 2024). Of course, the low confidence in AlphaFold predictions of Spo11 dimers could also reflect the fact that the Spo11 dimer interface is indeed weak compared to that of Top6A, or could be due to the absence of Spo11 dimers in structural databases.

Conservation analysis of the dimer interfaces of Spo11 and Top6A did not reveal obvious differences in their conservation score, suggesting that, despite the monomeric state of Spo11, the integrity of its dimerization surface remains under relatively high selective pressure (**Supplementary Figure 8B**). This selective pressure may balance the strength of the dimer interface itself: too strong could be toxic and too weak would lead to sterility.

### A Spo11-A236V mutation pokes within the active site across the dimer interface

In parallel to our site-directed mutagenesis approach, we performed a forward genetic screen in search for mutations that abolish meiotic DSB formation. The screen is based on the notion that DSB-defective mutants bypass the meiotic arrest of a *dmc1* mutant, which can be screened visually by auto-fluorescence of yeast spores (Hochwagen et al. 2005). From this screen, we identified five *spo11* alleles (P92V, A236V, G259R, G318E, C375Y), likely to be severe loss-of-function mutants. Purification attempts showed that most of the mutants were unstable, which accounts for their phenotype, except for Spo11-A236V.

Interestingly, A236 is highly conserved (**Supplementary Figure 3**) and AlphaFold modeling suggests that it is located at the dimer interface of Spo11 (**Figure 3B**). To characterize the Spo11-A236V mutant biochemically, we purified core complexes carrying the mutation and analyzed the DNA-binding activity of the complex by gel shift assay. On a short hairpin substrate containing a 2-nt 5′-overhang, the A236V mutant bound the substrate as efficiently as the wild type, indicating that end-binding is not affected (**Supplementary Figure 9**). Similarly, linear and branched substrates with 3′-overhangs were bound efficiently, and the assembly of Complex 2 was not affected, indicating that the Spo11-A236V mutation does not impact dimerization (**Figure 5A, B, C, Supplementary Figure 9**).

**Figure 5:**
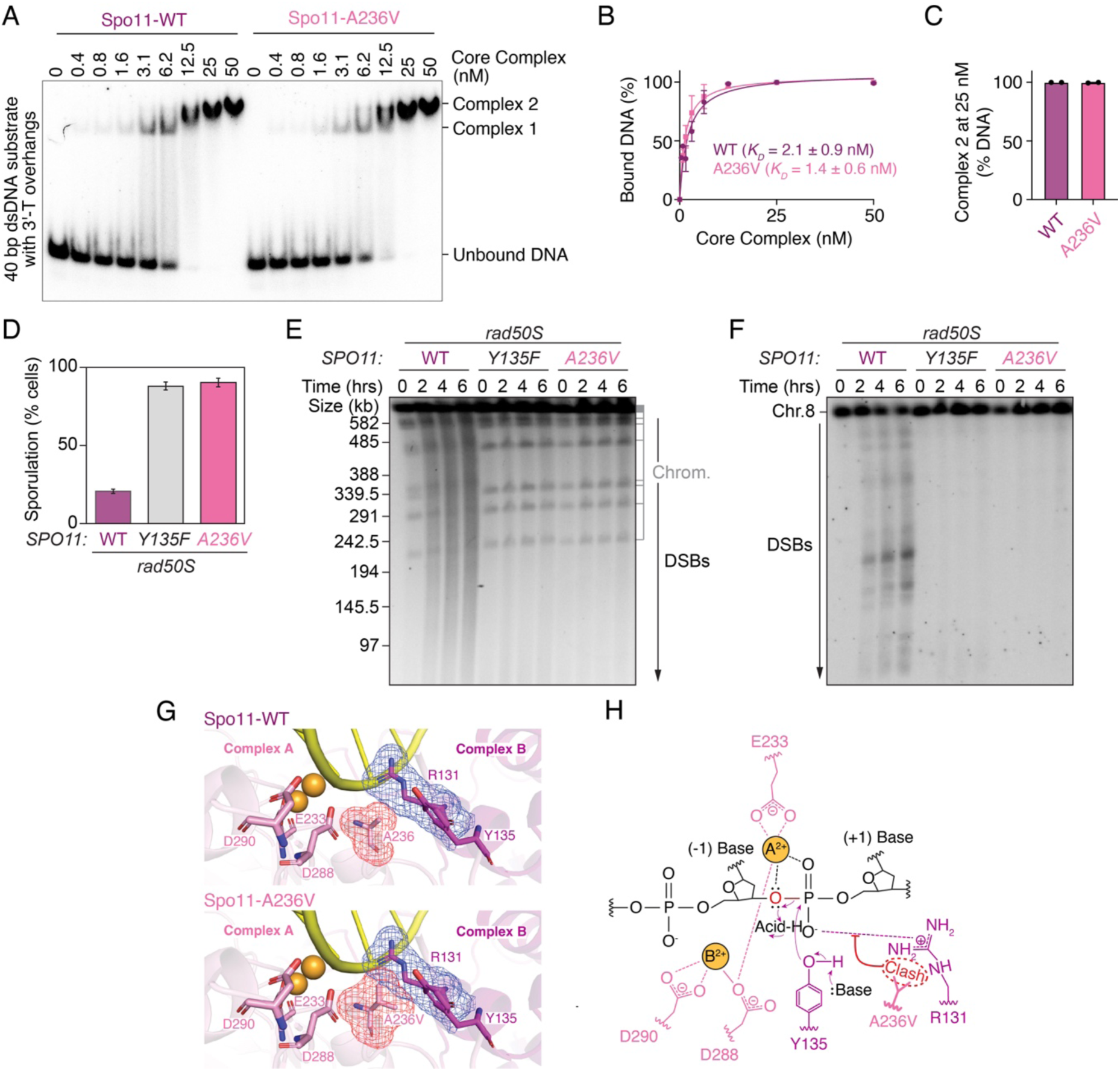
A Spo11-A236V mutation abolishes DNA cleavage via a steric clash across the dimer interface. **(A)** Gel shift analysis of DNA binding by wild-type and Spo11-A236V mutant core complexes to a 40 bp dsDNA substrate with 1-nt 3′-overhangs. **(B)** Quantification of the gel shift analysis of wild-type and Spo11-A236V mutant core complexes. **(C)** Quantification of Complex 2 at 25 nM core complexes (mean ± range of two independent experiments). **(D)** Sporulation frequency in wild-type, *spo11-Y135F* and *spo11-A236V* strains (*rad50S* background). **(E, F)** Pulse-field gel electrophoresis and Southern blot analysis of DSB formation in wild-type, *spo11-Y135F* and *spo11-A236V* mutant strains (*rad50S* background). **(G)** Modeling of the active site of Spo11-A236V core complexes. A236V and the active site residues (R131, Y135, E233, D288, D290) are shown as sticks, with metal-binding residues from the Toprim domain of Complex A (E233, D288, D290) colored in light pink and residues from the WH domain of Complex B (R131, Y135, A236, A236V) colored in magenta. The electron density map around A236V (red) and R131 (blue) are shown as mesh. In the AlphaFold3 model of the wild-type core complex active site (top), the β carbon (CB) of Spo11-A236 from Complex A lies 3.7 Å from the guanidinium terminal nitrogen 1 (NH1) and 3.4 Å from the δ carbon (CD) of R131 in Spo11 from Complex B. Given the van der Waals radii for carbon (1.70 Å) and nitrogen (1.55 Å), the expected non-bonded C···N distance is ∼3.25 Å. Both contacts are therefore greater than the expected non-bonded C···O distance, indicating no steric overlaps. In the AlphaFold3 model of the Spo11-A236V core complex active site (bottom), the γ2 carbon (CG2) of A236V in Spo11 from Complex A lies 2.3 Å from the NH1 and 2.6 Å from the CD of R131 in Spo11 from Complex B. Given the van der Waals radii for carbon (1.70 Å) and nitrogen (1.55 Å), the expected non-bonded C···N distance is ∼3.25 Å. Contacts are therefore ∼0.95 Å (CG2-NH1) and ∼0.65 Å (CG2-CD) shorter than expected, exceeding the 0.4 Å steric clash threshold used by MolProbity (Chen et al. 2010), and indicate significant steric overlaps. **(H)** The A236V mutation would perturb the two-metal-ion reaction mechanism of Spo11 core complex, based on the mechanism proposed for type IIA topoisomerases (Schmidt et al. 2010).

To verify the effect of the *spo11-A236V* mutation *in vivo*, we introduced the mutation in the context of yeast strains that harbor the *rad50S* allele, which prevents DSB repair and causes a sporulation defect. Similar to the catalytically inactive *spo11-Y135F* mutant, the *spo11-A236V* mutant restored sporulation in this background, indicative of a strong DSB defect (**Figure 5D**). Consistently, pulse-field gel electrophoresis and Southern blot analysis of chromosome VIII showed that the *spo11-A236V* mutant failed to produce any detectable DSBs (**Figure 5E, F**).

To understand what causes this phenotype, we modeled this mutation structurally. This revealed that the A236V allele would be expected to create a steric clash across the dimer interface, poking into active site residue R131 (**Figure 5G, H**), which has an essential catalytic function (Diaz et al. 2002). This accounts for the strong phenotype of the A236V mutation and provides further support for the close collaboration between two Spo11 subunits during catalysis.

### DNA binding by Rec102 is important for DSB formation

We sought to further investigate the contribution of Rec102 during DSB formation. Previous work showed that the B-subunit of Topo VI participates in substrate recognition and DNA bending through interactions between positive patches at the surface of Top6B and DNA (Wendorff and Berger 2018). At the time this work was initiated, no high-quality structural model was available for the Spo11 core complex (Claeys Bouuaert et al. 2021b), and the sequences of Top6B and Rec102 are too divergent to reliably model mutations based on Top6B (Robert et al. 2016). Hence, to test the possible importance of DNA binding by Rec102, we opted for an extensive site-directed mutagenesis screen of positively charged residues. We also mutated Rec104 residues given its close partnership with Rec102 (Claeys Bouuaert et al. 2021b).

We sought to identify mutations that reduce recombination based on a heteroallele recombination assay, but do not impact the interaction between Rec102 and Rec104 in a yeast two-hybrid assay. None of the eight Rec104 mutants tested had any significant effect on recombination (**Supplementary Figure 10**). Among the eighteen Rec102 mutants tested, we identified four alleles that abolished meiotic recombination (H171D, R197E, R199E/K201E, and R243E/R244E/R245E) (**Figure 6A**, left). However, all except one (R243E/R244E/R245E, RRR) also strongly disrupted the interaction with Rec104 (**Figure 6A**, right).

**Figure 6:**
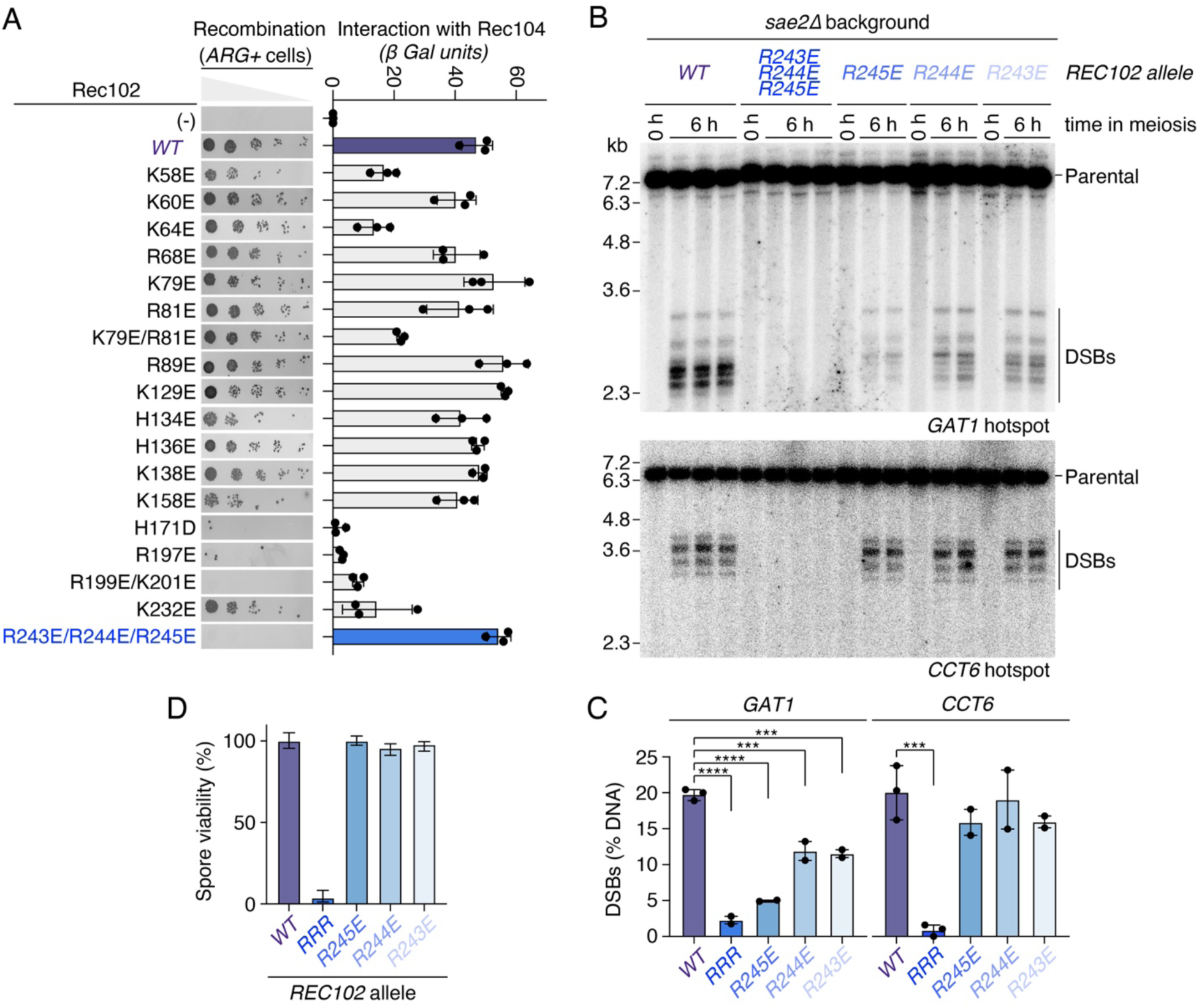
A positive patch in Rec102 is important for DSB formation. **(A)** Mutagenesis analysis of positively charged Rec102 residues on recombination established by *arg4* heteroallele recombination assay and Rec104 interaction established by yeast two-hybrid assay. **(B)** Southern blot analysis of DSB formation at the *GAT1* and *CCT6* hotspots in wild-type and Rec102 mutants in a *sae211* background. **(C)** Quantification of gels in panel B (mean ± range of 2-3 biological replicates per strain). **(D)** Spore viability analysis of wild-type and Rec102 mutant strains. n = 80 (WT), 136 (R243E), 160 (R244E), 136 (R245E), 136 (RRR) spores, respectively. Error bars are 95% confidence intervals of the proportion. and complexes with the Rec102-RRR mutation had >100-fold reduced DNA-binding activity (Figure 7B). Rec102 therefore contributes crucially to DNA binding by the Spo11 core complex.

To test the contribution of each residue of this motif, we mutated the three arginine residues individually or in combination at their endogenous locus. Southern blot analysis in a *sae2Δ* background indicated that all three mutants individually reduce DSB formation at the *GAT1* hotspots with the strongest reduction observed for the R245E mutant, while the triple mutant failed to produce detectable DSBs (**Figure 6B**). Similarly, no breaks were detected with the triple mutant at the *CCT6* hotspot, but the single mutants were indistinguishable from the wild type. Consistently, the spore viability of the triple mutant was 0-5%, while that of the single mutants was similar to the wild type (**Figure 6C**). To address whether these phenotypes are due to an impact of the mutations on DNA binding, we purified core complexes harboring the Rec102-R245E and Rec102-RRR mutations and analyzed their DNA-binding activities by gel shift assay (**Figure 7A**). Core complexes with the Rec102-R245E mutation had 20−50-fold reduced DNA-binding activity to the three substrates tested (dsDNA, HJ and hairpin substrate), While this work was underway, a cryoEM structure of the Spo11 core complex bound to a DNA end was solved (Yu et al. 2024). This structure that represents a post-cleavage state showed Rec102 residue R245 in contact with the DNA, 12 nt 3′ from the dyad axis. Our results show that this interaction is indeed important, and that Rec102 must engage DNA prior to catalysis, perhaps facilitating the bending of the DNA substrate. Consistently, the AlphaFold model of a pre-cleavage complex also places this positive patch in contact with DNA positioned in the major groove about one helix turn from the dyad axis (**Figure 7C**).

**Figure 7:**
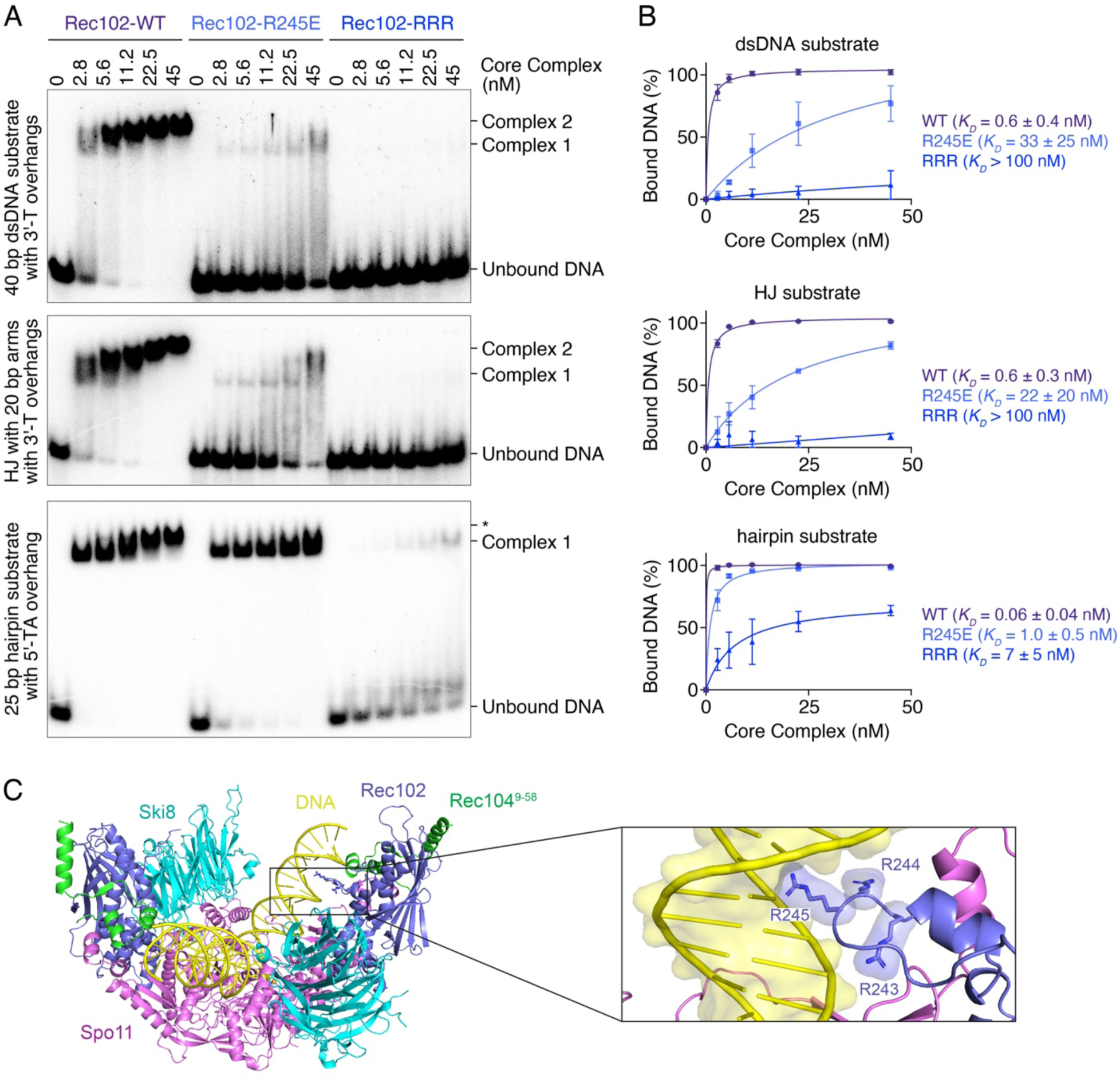
Rec102 directly contributes to DNA binding. **(A)** Gel shift analysis of the binding activity of Rec102 wild-type and mutant core complexes to DNA duplexes (40 bp dsDNA with 1-nt 3′-overhangs), Holliday junctions (HJ with 20 bp arms and 1-nt 3′-overhangs), or DNA ends (25 bp hairpin substrates with a 2-nt 5′-overhang). The band labeled * corresponds to monomeric complexes likely bound to opposite ends of the hairpin DNA substrate. **(B)** Quantification of the affinity of wild-type and mutant complexes for each substrate (mean ± range of two independent experiments). **(C)** Position of Rec102 residues R243, R244 and R245 in the AlphaFold model of the pre-cleavage Spo11 core complex.

### Rec102 contacts Ski8 in trans to facilitate dimer assembly

From the Rec102 mutants initially screened by heteroallele recombination assay, a few mutants appeared to have a mild recombination defect, including Rec102-K129E (**Figure 6A**). On the AlphaFold model, Rec102-K129 is not in contact with DNA, but is instead located on a surface that contacts Ski8 in *trans* across the dimerization plane (**Figure 8A**). The Rec102-Ski8 interface predicted by AlphaFold is composed of conserved residues that form somewhat hydrophobic patches of opposite electrostatic potential, with the Rec102 surface showing overall positive charge and the Ski8 surface overall negative charge (**Figure 8B-D**). This *trans* Rec102-Ski8 interaction was found in AlphaFold models of core complexes across ascomycetes (**Supplementary Figure 12**).

**Figure 8:**
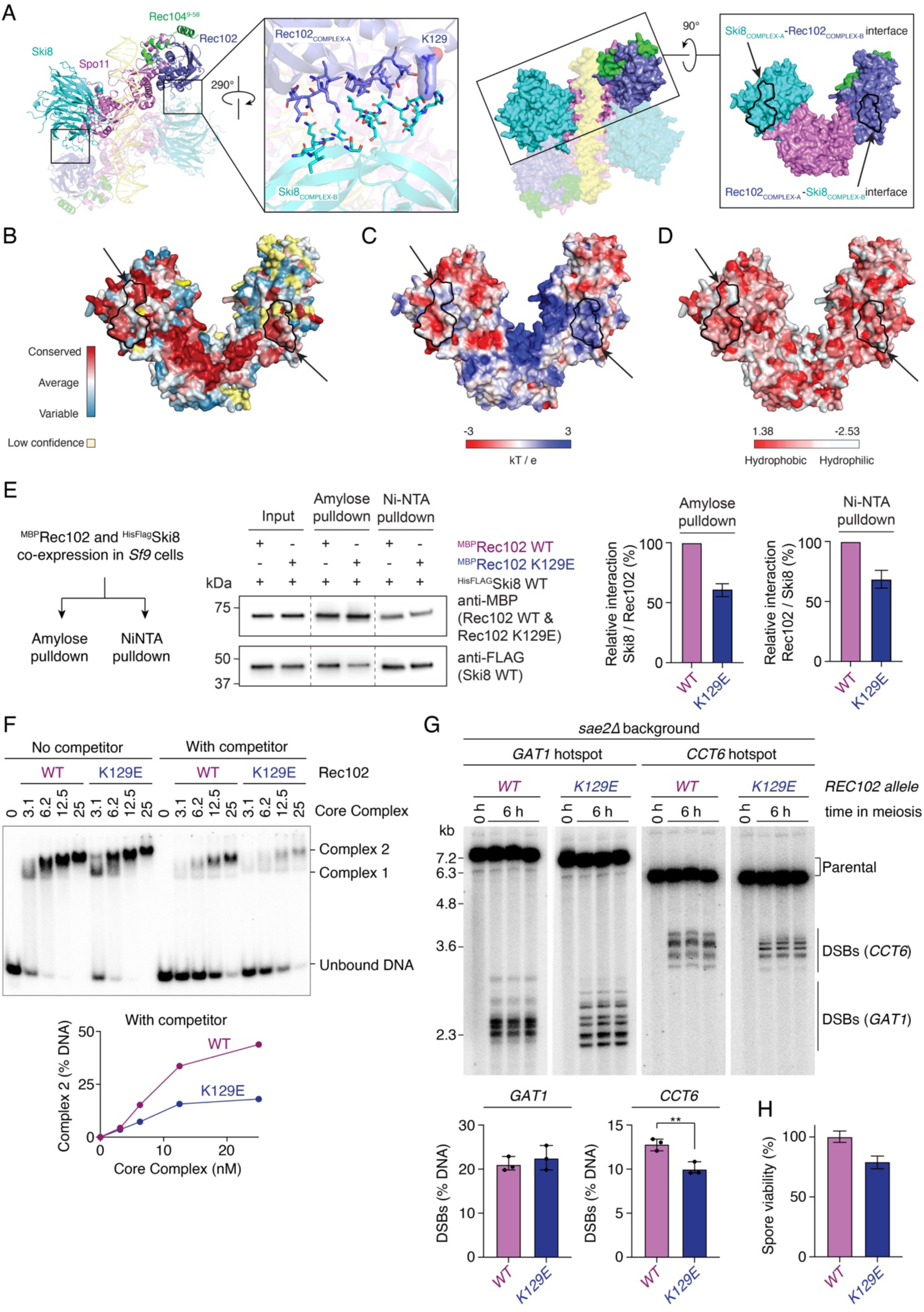
*Trans* contacts between Rec102 and Ski8 stimulate dimerization and DSB formation. **(A)** AlphaFold model of a dimeric Spo11 core complex bound to DNA. Complex B and DNA are shown semi-transparent. Middle-left, zoom of the predicted Rec102-Ski8 interface, with interface residues shown as sticks and the surface of K129 shown. Middle-right, surface representation of the model. Right, surface of a monomeric Spo11 core complex with predicted Ski8-Rec102 and Rec102-Ski8 interfaces outlined in black and indicated by arrows. **(B, C, D)** Conservation, electrostatic surface potential, and hydrophobicity of the dimerization surface with predicted Rec102-Ski8 interfaces outlined and indicated with arrows. **(E)** Pull-down analysis of MBP-tagged Rec102 wild-type and K129E mutant with HisFlag-tagged Ski8. **(F)** Gel shift analysis of DNA binding by wild-type and Rec102-K129E mutant core complexes on a radiolabeled 40 bp dsDNA substrate (0.5 nM) with or without 20 bp dsDNA competitor (25 nM). **(G)** Southern blot analysis of DSB formation at the *GAT1* and *CCT6* hotspots in wild-type and *rec102-K129E* mutant in a *sae211* background. **(H)** Spore viability analysis of wild-type and *rec102-K129E* mutant strains. n = 80 (WT) and 240 (K129E) spores, respectively. Error bars are 95% confidence intervals of the proportion.

In the monomeric core complex, Ski8 and Rec102 are bound at opposite ends of Spo11 and do not contact each other (Claeys Bouuaert et al. 2021b; Yu et al. 2024). Nevertheless, co-expression of MBP-tagged Rec102 with His-tagged Ski8 followed by NiNTA or amylose pull-down indicated that the proteins indeed interact *in vitro*, supporting the possibility that they interact in the context of a dimeric structure (**Figure 8E**). The Rec102-K129E mutation reduced this interaction by 30-40% (**Figure 8E**).

Next, we tested the impact of the Rec102-K129E mutation on dimerization of the core complex by gel shift assay. While the effect of the K129E mutation on the assembly of Complex 2 was subtle in our standard conditions, the addition of a fast-migrating (20 bp) competitor DNA substrate led to a marked reduction in Complex 2 with the K129E mutant, compared to the wild type (**Figure 8F**, **Supplementary Figure 11**). Hence, the Rec102-K129E mutation has a weak but detectable effect on dimerization of the core complex.

To test the effect of the mutation *in vivo*, we introduced the mutation at the endogenous *REC102* locus and characterized the effect of the mutation on DSB formation by Southern blot analysis at the *GAT1* and *CCT6* hotspots in a *sae2Δ* background. As compared to the wild type, the *rec102-K129E* mutant showed redistributed breaks at both hotspots, with a slight reduction of DSB levels detected at *CCT6* (**Figure 8G**). This led to a mild spore viability defect (79 %) (**Figure 8H**). Similar to the *spo11-S136A/V138A/E139A* dimerization mutant, the impact of the mutation on the position of cleavage sites is likely a consequence of reduced efficiency in the assembly of a cleavage-competent complex.

## Discussion

Here, we investigated the mechanism of Spo11 dimerization and the role of its accessory partners Ski8, Rec102, and Rec104. We show that Spo11 dimerizes transiently on DNA to form unstable complexes characterized by rapid subunit exchange. Despite the key role of dimerization in catalysis, the Spo11 dimerization surface is highly tolerant to mutations. Nevertheless, we identified a triple mutation (S136A/V138A/E139A) that significantly reduces dimerization on DNA and leads to reduced and redistributed DSBs *in vivo*, and identified another mutation (A236V) that abolishes DSB formation through occlusion of the active site across the dimer interface. Finally, we showed that Rec102 exerts an essential DNA-binding function and promotes dimerization through *trans* contacts with Ski8. This work yields new insights into the mechanisms whereby Spo11 and its partners collaborate to assemble pre-cleavage complexes.

### Spo11 dimerization

We previously showed that the *S. cerevisiae* core complex is monomeric in solution (Claeys Bouuaert et al. 2021b), which is also the case of the *M. musculus* (Oger and Claeys Bouuaert 2025b; Tang et al. 2025; Zheng et al. 2025) and *C. elegans* (Yeh et al. 2017) orthologs. This suggests that the monomeric state is a fundamental and conserved feature of Spo11 that may participate in preventing uncontrolled DSB formation. Indeed, we and others have shown that dimerization is a key factor limiting the DNA cleavage activity of mouse SPO11 *in vitro* (Oger and Claeys Bouuaert 2025b; Zheng et al. 2025). Given the essential role of dimerization for catalysis, we were surprised to find that most mutations at the predicted dimer interface of *S. cerevisiae* Spo11 have little impact on DSB formation *in vivo*. Nevertheless, *in vitro*, the mutations we analyzed (F203A/T239A/N243A, H246A/K247A, and S136A/V138A/E139A) all led to reduced assembly of dimeric complexes, indicating that the dimerization surface was indeed targeted. Hence, while the assembly of dimeric nucleoprotein complexes *in vitro* strongly relies on the Spo11 dimer interface, DNA cleavage *in vivo* does not, suggesting that additional mechanisms facilitate Spo11 dimerization *in vivo*. Likely mechanisms include interactions with Rec114 and Mre11, which are constitutive dimers themselves (Williams et al. 2008; Daccache et al. 2023) and directly bind Rec102-Rec104 and Spo11, respectively (Maleki et al. 2007; Claeys Bouuaert et al. 2021a; Aithal et al. 2024).

### A new role for Rec102 and Ski8 in dimerization

Based on AlphaFold modeling, we found that Rec102 contacts Ski8 across the dimerization plane, suggesting that this interaction supports dimerization. Indeed, we found that the Rec102-K129E mutation led to a mild reduction in Rec102-Ski8 interactions and in the assembly of dimeric nucleoprotein complexes *in vitro*, and concomitant reduction and redistribution of DSBs *in vivo*.

We note that the Rec102-K129E allele was not designed to abolish the interaction with Ski8 and that the cause of its meiotic phenotype was uncovered *a posteriori* through AlphaFold modeling. Targeted mutagenesis of this interface is likely to reveal more severe mutants that will be better suited to probe the physiological importance of this interaction. Interestingly, while several mutations of the Spo11 dimer interface (F203A/T239A/N243A and H246A/K247A) that clearly disrupted the assembly of dimeric complexes *in vitro* had no discernable impact on DSB formation *in vivo*, the Rec102-K129E mutation that disrupted the assembly of the dimeric complex more subtly (evident only when we challenged the complexes with competitor DNA), caused a clearer effect on DSB formation *in vivo*. This suggests that the *trans* Rec102-Ski8 interaction that we describe is comparatively more important for DSB formation than Spo11-Spo11 interactions.

What is the purpose of this Rec102-Ski8 interaction? Why does Spo11 rely on partners for dimerization? We propose that delegating that responsibility to accessory partners allows for a finer control of Spo11 activity, including through non-linearity effects of protein expression. For instance, Ski8 overexpression may inhibit Spo11 dimerization by occluding the low-affinity binding site of Rec102, and *vice versa*. Similarly, any effect of Spo11 overexpression may be buffered if not incorporated within *bona fide* core complexes.

While Rec102 is an ancient partner of Spo11 (Robert et al. 2016), Ski8 is likely a later addition to the meiotic DSB core complex. Indeed, Ski8 is required for DSB formation in *S. cerevisiae*, *S. macrospora* and *S. pombe*, but not in *A. thaliana* (Tesse et al. 2003; Jolivet et al. 2006; Steiner et al. 2010). Consistently, AlphaFold modeling suggested that Ski8 participates in dimerization of the core complex throughout ascomycetes.

### The role of Rec102 in DNA binding

In addition to this new role in dimerization of the core complex, we also show that Rec102 exerts an essential DNA-binding function. A cryo-EM structure obtained while this work was ongoing showed that Rec102 binds DNA in the context of a post-cleavage complex (Yu et al. 2024). Here, we show that this interaction precedes catalysis, that it enhances the affinity of the core complex to all the DNA substrates tested by several orders of magnitude, and that it is essential for DSB formation.

Previous work had shown that Rec102’s distant relative, Top6B, participates in substrate recognition and DNA bending to coordinate the capture of two DNA duplexes with ATP-driven dimerization of its GHKL domain and DNA cleavage by the Top6A subunit (Wendorff and Berger 2018). Although the sequences of Top6B and Rec102 evolved almost beyond recognition, our findings suggest that the DNA-binding function of the ancestral B-type subunit was conserved in Rec102. However, whether Rec102 participates in substrate selection and DNA bending *per se* is unclear. Indeed, the strongly reduced DNA-binding activity of the Rec102-R245E and Rec102-RRR mutants prevented us from addressing that question.

A comparison of the *in vitro* and *in vivo* phenotypes of these mutants provides further evidence of the robustness of the DSB machinery. Indeed, while the Rec102-R245E mutation caused a 20 to 50-fold reduction in the affinity of the core complex for DNA, the frequency of DSB formation as assayed by Southern blotting in a *sae2Δ* background was only reduced about 4-fold at *GAT1* and hardly reduced at all at *CCT6*, leading to no spore viability defect. Nevertheless, the triple Rec102-RRR mutation that further reduced DNA binding at least 5-fold abrogated DSB formation, causing ∼100% spore lethality. Hence, the contribution of Rec102 to the DNA-binding activity of the core complex is essential, even though the cell can tolerate a significant reduction in affinity.

### Does the Spo11 core complex capture two duplexes?

Our previous AFM analyses using plasmid DNA substrates indicated that the core complex often binds at the junction between two DNA duplexes, a property shared with other type II topoisomerases (Alonso-Sarduy et al. 2011; Timsit 2011; Wendorff and Berger 2018). Here, we found that, like Topo VI (Wendorff and Berger 2018), the core complex efficiently binds four-way branched DNA structures that mimic two juxtaposed DNA duplexes, and it efficiently dimerizes on this substrate through Spo11-Spo11 and Ski8-Rec102 interactions. Although a detailed understanding of this assembly will require high-resolution structural analyses, we note that the dimensions of the arms (20 bp) in our substrate are insufficient to accommodate a core complex dimer, and therefore the complex must have bound the junction itself.

The relevance of this junction-binding activity can be questioned, given that the biological function of Spo11 does not require strand passage. Nevertheless, the hypothesis that this activity is significant is supported by previous work showing that long palindromic repeats inserted within the *S. cerevisiae* genome create meiotic DSB hotspots (Nasar et al. 2000). A straightforward interpretation is that these sequences extrude as cruciforms, which are structurally equivalent to the Holliday Junctions we analyzed. Since this creates a hotspot in yeast, it suggests either that the cruciforms are bound preferentially by Spo11, or that they somehow activate cleavage by mimicking complexes with two bound DNA duplexes.

If this is indeed the case and Spo11 must trap two duplexes prior to cleavage, the question arises as to what this additional duplex corresponds to at natural hotspots. An intriguing possibility is that the core complex would capture both sister chromatids. This would provide an effective mechanism to prevent Spo11 from cleaving both sisters (Zhang et al. 2011).

Nevertheless, other scenarios are conceivable, and the requirement for binding two duplexes may not be absolute. Indeed, we have shown that mouse SPO11 can cleave DNA by itself *in vitro* (Oger and Claeys Bouuaert 2025b). In the absence of a B-type subunit, a SPO11 dimer could not capture two DNA duplexes, excluding the possibility that this is an essential prerequisite for cleavage. Nevertheless, the scenario could be different *in vivo*, where TOP6BL and other partners are present.

### The enigmatic function of Rec104

Finally, while our structure-function analysis brought new insights into how Spo11 dimerizes and uncovered new functions for Rec102 and Ski8, the role of Rec104 remains somewhat enigmatic. Rec104 is essential for the integrity of the core complex (Claeys Bouuaert et al. 2021b) and directly participates in protein interactions with Rec114 (Maleki et al. 2007; Claeys Bouuaert et al. 2021a). The N-terminal 60 amino acids interact with Rec102 and the C-terminal part is probably poorly structured (Yu et al. 2024).

Rec104 is phosphorylated during meiosis (Kee et al. 2004), but the functional importance of this phosphorylation has not been established. Compared to other members of the complex, Rec104 is poorly conserved. Nevertheless, we uncovered that the C-terminal extremity of Rec104 contains a highly conserved motif that matches a putative site for CDK phosphorylation. CDK is known to phosphorylate Mer2, which is essential for DSB formation (Henderson et al. 2006). It will be interesting to address whether CDK also phosphorylates Rec104, and whether this has physiological consequences.

## Materials and Methods

### Preparation of plasmid vectors

Plasmids are listed in **Supplementary Table 2**. Oligonucleotides are listed in **Supplementary Table 3**. pFastBac1-based vectors to express wild-type Spo11^HisFlag^ (pCCB592), Ski8 (pCCB587), Ski8^HisFlag^ (pCCB615), Rec102 (pCCB588), ^MBP^Rec102 (pCCB617) and Rec104 (pCCB589) were previously described (Claeys Bouuaert et al. 2021b). A Tev recognition site was added to plasmid pCCB617 between the coding sequences of MBP and Rec102 by PCR amplification with primers ab001 & ab002 followed by blunt-end ligation to yield pHAB001 (^MBP-tev^Rec102). The coding sequence of mVenus was amplified with primers ab133 & ab134 and cloned into the product of pHAB001 amplification with primers ab135 & ab136 by Gibson assembly to yield pHAB054 (^MBP-tev-YFP^Rec102).

#### Spo11 mutagenesis

For the heteroallele recombination assay and yeast strain constructions, the template (pCCB767) contained the *SPO11-HisFlag::HphMX6* cassette cloned into pRS314. For baculovirus production, the template was pCCB592 (Spo11^HisFlag^ in pFastBac1) (Claeys Bouuaert et al. 2021b). Spo11 mutants were generated by site-directed mutagenesis using the following primers: Spo11-K126A/N127A (cb991 & cb992), Spo11-K206A/P207A (cb995 & cb996), Spo11-T239A/K240A (cb997 & cb998), Spo11-F203A (ab228 & ab229), Spo11-F203A/K206A/P207A (ab230 & ab231), Spo11-S136A/V138A/E139A (ab234 & ab235), Spo11-T239A/N243A (ab216 & ab217), Spo11-K240A (ab275 & ab276), Spo11-H246A/K247A (ab293 & ab294), Spo11-A236V (ab253 & ab254).

#### Rec102 mutagenesis

For yeast-two-hybrid and heteroallele recombination assays, the template was pSK282 (LexA-Rec102 plasmid) (Maleki et al. 2007). For baculovirus production, the template was pHAB001 (^MBP-tev^Rec102 in pFastBac1). For yeast strain constructions, the template was pCCB758 (Rec102 followed by the NatMX6 cassette). Rec102 mutants were generated by site-directed mutagenesis using the following primers: Rec102-K58E (ab067 & ab068), Rec102-K60E (ab065 & ab066), Rec102-K64E (ab063 & ab064), Rec102-R68E (ab061 & ab062), Rec102-K79E/R81E (ab033 & ab034), Rec102-K79E (ab035 & ab036), Rec102-R81E (ab037 & ab038), Rec102-R89E (ab039 & ab040), Rec102-K129E (ab041 & ab042), Rec102-H134A (ab043 & ab044), Rec102-H136A (ab045 & ab046), Rec102-K138E (ab047 & ab048), Rec102-K158E (ab049 & ab050), Rec102-H171D (ab051 & ab052), Rec102-R197E (ab053 & ab054), Rec102-R199E/K201E (ab055 & ab056), Rec102-K232E (ab057 & ab058), Rec102-R243E/R244E/R245E (ab059 & ab060), Rec102-R243E (ab299 & ab300), Rec102-R244E (ab301 & ab302), Rec102-R245E (ab218 & ab219).

#### Rec104 mutagenesis

For yeast-two-hybrid and heteroallele recombination assays, the template was pSK293 (Rec104-LexA) (Arora et al. 2004). For baculovirus production, the template was pCCB589 (Rec104 in pFastBac1) (Claeys Bouuaert et al. 2021b). Rec104 mutants generated by site-directed mutagenesis are: Rec104-H19D (ab017 & ab018), Rec104-K39E/R40E (ab019 & ab020), Rec104-K48E (ab021 & ab022), Rec104-K65E/R66E (ab023 & ab024), Rec104-K78E/K79E (ab025 & ab026), Rec104-K97E (ab027 & ab028), Rec104-R122E (ab029 & ab030), Rec104-R147E (ab031 & ab032).

### Protein expression and purification

Core complex proteins were expressed in *Spodoptera frugiperda* Sf9 cells using the Bac-to-Bac baculovirus expression system (Invitrogen^TM^) and purified essentially as described (Claeys Bouuaert et al. 2021b). Spo11^HisFlag^:^MBP^Rec102:Rec104:Ski8 complexes used viruses produced from vectors pCCB592, pHAB001, pCCB589, and pCCB587. Other complexes were prepared with appropriate combination of viruses (see list of vectors in **Supplementary Table 2**). Cultures of Sf9 cells (1 L containing 2.10^6^ cells/mL) were co-infected with each virus at a multiplicity of infection of 2. Cells were agitated at 100 rpm at 27 °C for 2.5 days, then harvested and washed twice with ice-cold phosphate-buffered saline (1X PBS). The cell pellet was either stored at - 80 °C or used immediately. All the purification steps were carried out at 0-4 °C. Cells were resuspended in a buffer containing 25 mM HEPES-NaOH pH 7.5, 20 mM imidazole, 0.1 mM DTT, 0.5 mM EDTA supplemented with 1× Complete protease inhibitor tablet (Roche) and 0.3 mM PMSF. Cells were lysed by osmotic shock by the addition of glycerol (final concentration of 10% v/v) + NaCl (final concentration of 500 mM). Cell extracts were cleared by ultracentrifugation (40 krpm) for 40 minutes and the supernatant was gently agitated for 1-3 hours with pre-equilibrated NiNTA agarose resin (Qiagen). Resin was collected by mild centrifugation (300 g) before being deposited into a disposable column for additional washing steps. Upon extensive washes with Nickel buffer (25 mM HEPES-NaOH pH 7.5, 20 mM imidazole, 500 mM NaCl, 0.1 mM DTT, 0.5 mM EDTA, 10% glycerol), protein complexes were eluted with buffer containing 500 mM imidazole. Elution samples were analyzed by SDS-PAGE and fractions containing proteins were pooled and loaded on a disposable column containing pre-equilibrated Amylose resin (NEB). The resin was washed extensively with Amylose buffer (25 mM HEPES-NaOH pH 7.5, 500 mM NaCl, 1 mM DTT, 2 mM EDTA, 10% glycerol), proteins were eluted with Amylose buffer supplemented with 10 mM of maltose, and analyzed by SDS-PAGE. Fractions containing proteins were pooled and separated by size-exclusion chromatography using either Superdex200 10/300 GL Increase or Superose6 10/300 GL Increase (Cytiva) columns. The peak of interest was collected, the complex was concentrated with Amicon centrifugal tubes (Sigma-Merck), aliquots were flash frozen in liquid N2 and stored at -80 °C.

### Pulldown assay

For pulldown analyses, wild-type or mutant ^MBP^Rec102 was co-expressed with Ski8^HisFlag^ in 50 mL Sf9 cells, resuspended in 1 mL buffer containing 25 mM HEPES 7.5, 500 mM NaCl, protease inhibitors (Roche Complete tablet + 0.5 mM PMSF), with either 20 mM imidazole, 0.2 mM DTT (for NiNTA affinity) or 1 mM DTT, 2 mM EDTA (for amylose affinity). Cells were lyzed by sonication, extracts cleared by centrifugation, and proteins were pulled down by NiNTA or amylose affinity. Upon extensive washes, proteins were eluted, separated by SDS-PAGE and analyzed by anti-His or anti-MBP Western blotting. To quantify the interaction between Ski8 and Rec102 variants, Ski8 signal intensity from the pull-down eluates was first normalized to the amount of Rec102 captured in the same pull-down. To correct for differences in Rec102 expression levels between conditions, these values were further normalized to Rec102 signal in input lysates. Band intensities were quantified using ImageJ, and background was subtracted.

### DNA substrates preparation

DNA substrates were generated by annealing complementary oligonucleotides (sequences in **Supplementary Table 3**). Oligonucleotides were mixed in equimolar concentrations in STE buffer (100 mM Tris pH 8, 1 mM EDTA, 100 mM NaCl) and annealed by heating and slow cooling in a thermocycler with the following program: 98 °C for 3 minutes, 75 °C for 1 hour, 65 °C for 1 hour, 37 °C for 30 minutes and 25 °C for 10 minutes. The 40 bp dsDNA substrate with 3′-T overhangs was prepared by annealing primers ab203 and ab204, the HJ substrate with 20 bp arms and 3′-T overhangs was prepared with primers ab185, ab186, ab187, ab188. The hairpin substrate with 5′-TA overhang was prepared by self-annealing of primer cb957. The 20 bp and 80 bp dsDNA substrates used as fast-migrating and slow-migrating competitors were made by annealing primers ab175, ab176, and cb95, cb100, respectively. Unlabeled substrates were gel purified on a native 10% TBE-polyacrylamide gel.

For 5’-end labeling with ^32^P, 5 pmol of cold substrates were incubated with 2 μL (approximately 20 μCi) of [γ-^32^P]-ATP (Revvity, 3000 Ci/mmol 10 mCi/ml). Upon labeling, excess of radionucleotides were removed by Quick Spin DNA column (Cytiva^©^) and substrates were gel-purified on a native 10% TBE-polyacrylamide gel, migrated for 1.5 hours at 150 V at 4 °C. To reveal radiolabeled substrates, X-ray films were laid on gels and kept in the dark before development and revelation. Bands were excised, the DNA was eluted in STE overnight at 4 °C, then DNA was ethanol precipitated, resuspended in 100 μL of STE and stored at −20 °C until use.

### Electrophoretic mobility shift assays

DNA-binding reactions were performed in 20 μL reactions containing 25 mM Tris-HCl pH 7.5, 10% glycerol, 100 mM NaCl, 5 mM MgCl2 and 1 mg/mL BSA with 0.2-1 nM of DNA substrates and the indicated amount of Spo11 core complexes. Unless specified, reactions were performed with core complexes that had an MBP tag fused to the N-terminus of Rec102 and a HisFlag tag fused to the C-terminus of Spo11. Some reactions also included cold DNA competitors in excess (25 nM, or as indicated). Samples were then incubated at 30 °C for 30 min before loading on native 5% Tris-Acetate-polyacrylamide gels (80:1 acrylamide:bisacrylamide) supplemented with 0.5 mM MgCl2, and complexes were separated at 180 V for 2.5 hours. Gels were dried at 80 °C for 30-60 minutes and exposed to autoradiography plates. Screens were revealed by phosphorimaging using a Typhoon scanner (Amersham), images were quantified with Fiji (ImageJ), and data was analyzed with Prism.

### Yeast targeting vectors and strain construction

Yeast strains are listed in **Supplementary Table 4**. For yeast strains expressing Spo11 or Rec102 mutants, the endogenous *SPO11* and *REC102* loci were replaced with mutant constructs marked by HphMX6 and NatMX6 resistance cassettes, respectively, located about 50 bp downstream of the coding sequence. For Spo11 mutants, vectors based on the pCCB767 plasmid were digested with SphI. For Rec102, vectors based on the pCCB758 plasmid were digested with EcoRI. DNA fragments were gel purified and used to transform the CBY006 yeast strain by lithium-acetate transformation. Transformants were selected on YPD plates containing the appropriate antibiotic, re-streaked on selective plates and genomic DNA of individual clones was isolated. The strains were first genotyped by PCR (primers cb1149 & cb1150 for *SPO11* and primers cb1273 & cb1274 for *REC102*), then the presence of the mutation was confirmed by sequencing. The *spo11-A236V* mutant was derived by EMS mutagenesis and confirmed by Sanger sequencing. Confirmed positive clones were saved, stored at -80 °C, and derivative strains were obtained by mating and tetrad dissections.

### Yeast-two-hybrid assay

Empty and wild-type vectors used for Y2H assays are: pSK276 (Gal4AD empty vector), pSK272 (LexA empty vector), pSK282 (LexA-Rec102), pSK293 (Rec104-LexA), pSK302 (GalAD-Rec102), and pSK310 (Rec104-Gal4AD) (Arora et al. 2004; Maleki et al. 2007). Mutant vectors were constructed as described above and are listed in **Supplementary Table 2**.

Haploid cells (CBY52 and CBY53) were transformed with yeast-two-hybrid vectors containing either *TRP1* (LexA fusions) or *LEU2* (Gal4 fusions) markers, plated on appropriate selective media (SC-W or SC-L) and incubated at 30 °C. After 2 to 3 days, cells were mated with strains of the opposite mating type and diploids were selected on double dropout (SC-WL) media. After 2 days, mated strains were streaked on SC-WL to get single colonies. After 2 or 3 days, single colonies were cultured in liquid selective medium supplemented with 2% glucose at 30 °C overnight. Cultures were then diluted into fresh liquid selective medium (SC-WL) supplemented with 2% galactose and 1% raffinose and incubated at 30°C. After 4 hours, cells were harvested and lyzed, and β-galactosidase activity was measured from ortho-nitrophenyl-β-galactopyranoside (ONPG) hydrolysis using standard protocols (Clontech laboratories).

### Heteroallele recombination assay

Heteroallele recombination assays were performed in diploid strains that carried the *arg4-Bgl* and *arg4-Nsp* alleles (Nicolas et al. 1989) (see **Supplementary Table 4**). Strains were diploids of CBY312 and CBY313 (for Rec102), CBY1111 and CBY1112 (for Spo11), and CBY1109 and CBY1110 (for Rec104). Haploid cells were first mated on YPD plates, then streaked to obtain single colonies. After two days at 30 °C, a diploid colony was incubated overnight in liquid YPD, then transformed with appropriate wild-type or mutant complementation plasmids (see **Supplementary Table 2**). Transformants were isolated on selective media (SC-W), single colonies were then cultured in selective media with vigorous shaking for 1.5 days. Cells were harvested, washed twice with H2O before meiotic induction in 2% potassium acetate pH 5.5. After 16 hours in meiosis, serial dilutions were spotted on plates lacking arginine to measure the frequency of Arg+ recombinants, and on YPD to measure colony forming units.

### Spore viability

Haploid cells were mated on YPD, then were transferred into 2-3 mL of 2% potassium acetate pH 5.5. Two days after meiotic induction, tetrads were dissected on YPD plates, and the frequency of colony forming units was calculated after 2 days at 30 °C.

### Southern blot analysis

Meiotic DSB analysis by Southern blotting was performed as previously described (Murakami et al. 2009). In brief, synchronized cultures undergoing meiosis were collected at the indicated time points. After DNA isolation, 1 µg of genomic DNA was digested by PstI-HF (NEB-R3140S) and separated on a 1% TBE-agarose gel (Lonza, seaKem LE agarose -50004) for 16 hours at 75 Volts. DNA was transferred to Amersham™ Hybond™-N+ nylon membranes (Cytiva-RPN303B) by vacuum transfer, hybridized with *GAT1* probe (amplified with primers: 5′-CGCGCTTCACATAATGCTTCTGG, 5’-TTCAGATTCAACCAATCCAGGCTC), or *SLY1* probe for *CCT6*: (amplified with primers 5′-GCGTCCCGCAAGGACATTAG, 5′-TTGTGGCTAATGGTTTTGCGGTG) and developed by autoradiography.

### Genetic screen for DSB-defective mutants

The *spo11-A236V* mutation was isolated from a forward genetic screen for mutations that bypass the sporulation defect caused by loss of *DMC1*. The details and full list of mutants resulting from this screen will be published elsewhere. Briefly, haploid *dmc1Δ spo13Δ* strains carrying both endogenous *MATa* and transgenic *MATα* mating type information were mutagenized with low levels of ethyl methanesulfonate (EMS). Individual mutant colonies were patched on YPD before inducing them to undergo sporulation on minimal sporulation medium (2% potassium acetate). Sporulation was monitored over 1-3 days at 30°C using a handheld UV lamp to assay for the expression of dityrosine in the spore wall (Hochwagen et al. 2005). Spore formation of UV-positive clones was confirmed under the microscope. To identify the mutated gene, bypass mutants were cured of the *MATα* transgene and crossed to a complementation panel of known bypass mutants before testing for non-complementation of the sporulation phenotype. The location of the causative mutation was determined by Sanger sequencing of the entire non-complementing gene. Sequencing of the *spo11-A236V* mutant revealed no other mutations in the *SPO11* open reading frame.

### Pulse-field gel electrophoresis and Southern blotting

Pulsed-field gel electrophoresis and Southern blotting for chromosome 8 was performed as described previously (Blitzblau et al. 2007). Briefly, cells were induced to undergo synchronous meiosis and samples were collected and inactivated with sodium azide at the indicated time points. Cells were embedded in agar plugs before treatment with zymolyase and proteinase K. Chromosomes were separated in 1% Seakem LE agarose in 0.5X TBE using a 5s-45s gradient for 32h at 5.4V/cm. The gel was stained with ethidium bromide and blotted onto a GeneScreen membrane (Revvity) using alkaline transfer. The membrane was probed for *CBP2* which is located near the left end of chromosome 8 (Blitzblau et al. 2007). Experiments were repeated as 2-4 biological replicates with consistent results.

### Bioinformatic analysis

Multiple sequence alignments of *S. cerevisiae* core complex proteins were generated using the sequences of one representative species of each of the 12 most well-accepted genera in *Saccharomycetaceae*. Sequences were aligned with MAFFT v7.511 using L-INS-i parameters for Spo11 and Ski8, and FFT-INS-i parameters of Rec102 and Rec104 (Katoh and Standley 2013). Multiple sequence alignments were visualized in Jalview with ClustalX coloring at 10% identity threshold (Waterhouse et al. 2009). Evolutionary conservation scores for each residue were calculated using ConSurf with *S. cerevisiae* Spo11 core complex proteins as query sequences (Ashkenazy et al. 2016). Site-specific evolutionary rates were calculated using the Bayesian inference method Rate4Site and the WAG substitution model (Pupko et al. 2002). Positions with <6 ungapped sequences or wide credibility intervals (≥4 Consurf color grades) were labeled ‘low confidence’ and not interpreted. Residues were labeled interface residues if the decrease in their Accessible Surface Area (ASA) upon complexation was larger than 1 Å^2^ (Jones and Thornton 1997).

BLAST searches for Rec102 homologs in *Nakaseomyces glabratus* (XP_003350816.1) identified sequences that lack one of the conserved β-strands and the characteristic WKxY motif (Robert et al. 2016). Inspection of the genomic locus suggests that a splice site is misannotated; correcting this results in a longer second exon, adding the missing sequence. We used the new sequence to model the structure using AlphaFold3.

## End Matter

### Author Contributions and Notes

H.A.B. designed mutants, performed RTG and Y2H experiments, purified proteins, performed gel shift assays, and generated yeast strains; M.S. performed Southern blot experiments; J.U.A. performed Rec102-Ski8 pulldowns, AlphaFold modeling, and bioinformatic analyses; V.V.S. performed the genetic screen for DSB mutants and D.D. performed *in vivo* analyses of the Spo11-A236V mutant under the supervision of A.H.; C.C.B. designed the research, secured funding, supervised the research, performed gel shift assays and analyzed data. C.C.B. wrote the paper with input from all authors. The authors declare no conflict of interest.

## Acknowledgments

We thank Manon Lenoir and Cédric Oger for comments on the manuscript. We also acknowledge the Core at the New York University Center for Genomics and Systems Biology for assistance with instrumentation. This work was supported by the European Research Council under the European Union’s Horizon 2020 research and innovation program (ERC grant agreement 802525 to CCB), and the Fonds National de la Recherche Scientifique (MIS-Ulysse grant F.6002.20 and PDR grant T.0031.22 to CCB). CCB is a FNRS research associate. This work was supported in part by the National Institutes of Health (grant R35GM148223 to AH).

## Supplementary Files

### Supplementary Figures

**Supplementary Figure 1:**
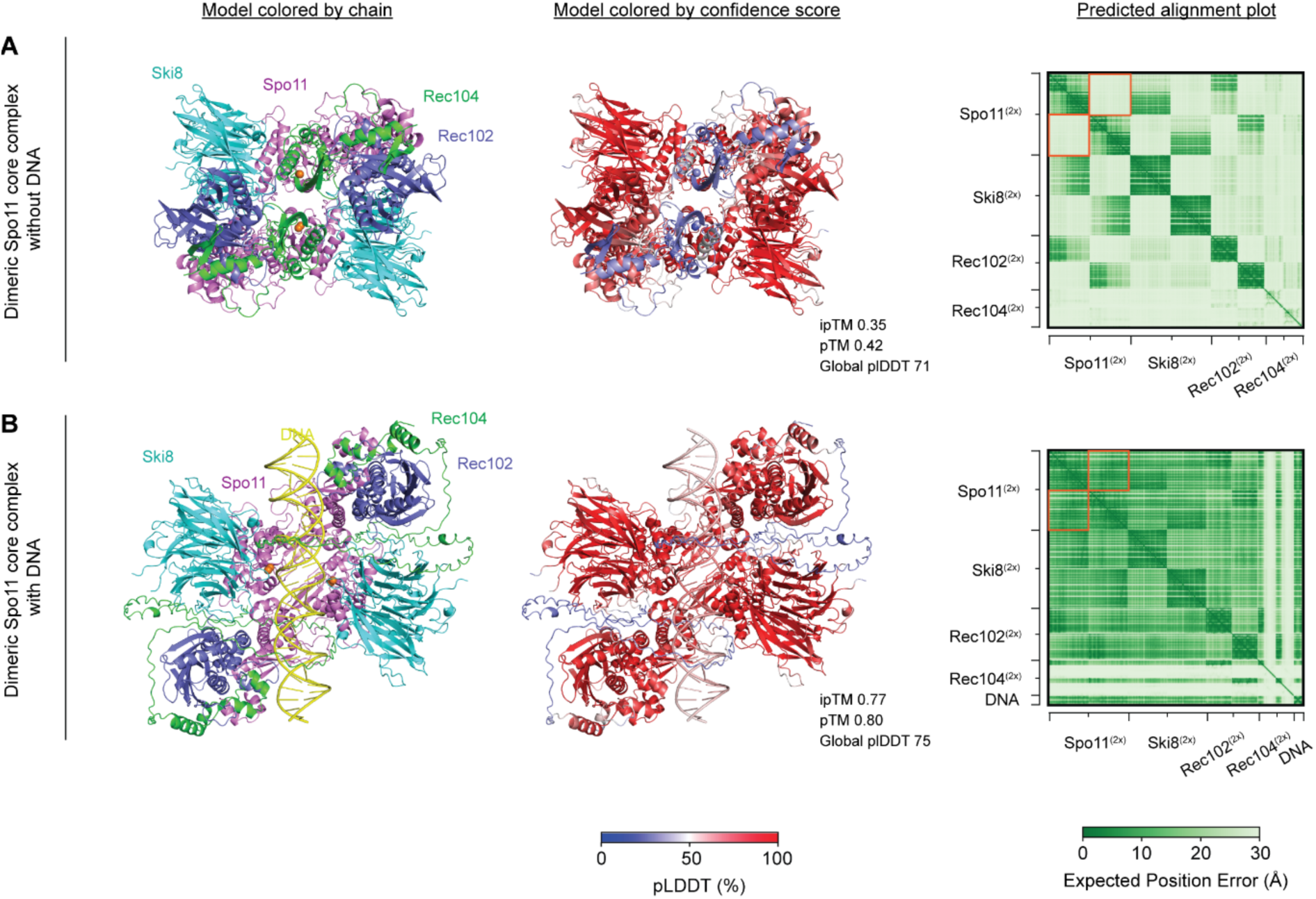
Quality assessment of structural models of dimeric Spo11 core complexes with and without DNA. AlphaFold3 models of 2:2:2:2 Spo11:Ski8:Rec102:Rec104 dimeric core complexes without (**A**) and with (**B**) a 40 bp duplex DNA substrate (5’-CGCACACACATCACACACCGCGGTGTGTGATGTGTGTGCG-3’). Left: AlphaFold3 models colored by chain. Spo11 (violet), Ski8 (cyan), Rec102 (blue), Rec104 (green), DNA (yellow) and Mg^2+^ ions (orange). Middle: AlphaFold3 models colored by confidence score. Left, predicted alignment error plot. In both models, the overall structure of Rec104 is predicted with low confidence, while those of Spo11, Ski8 and Rec102 are predicted with high confidence. The confidence in the relative position of Spo11 from Complex1 with Spo11 from Complex2 is indicated by the orange squares on the predicted alignment error plots. (**A**) In the absence of DNA, the relative position of the two Spo11:Ski8:Rec102:Rec104 heterotetramers is predicted with low confidence, and AlphaFold3 predicts an aberrant Spo11 dimer interface. (**B**) In the presence of DNA, the relative position of the two copies of the core complex is predicted with high confidence, and AlphaFold3 predicts a reliable Spo11 dimer interface.

**Supplementary Figure 2.**
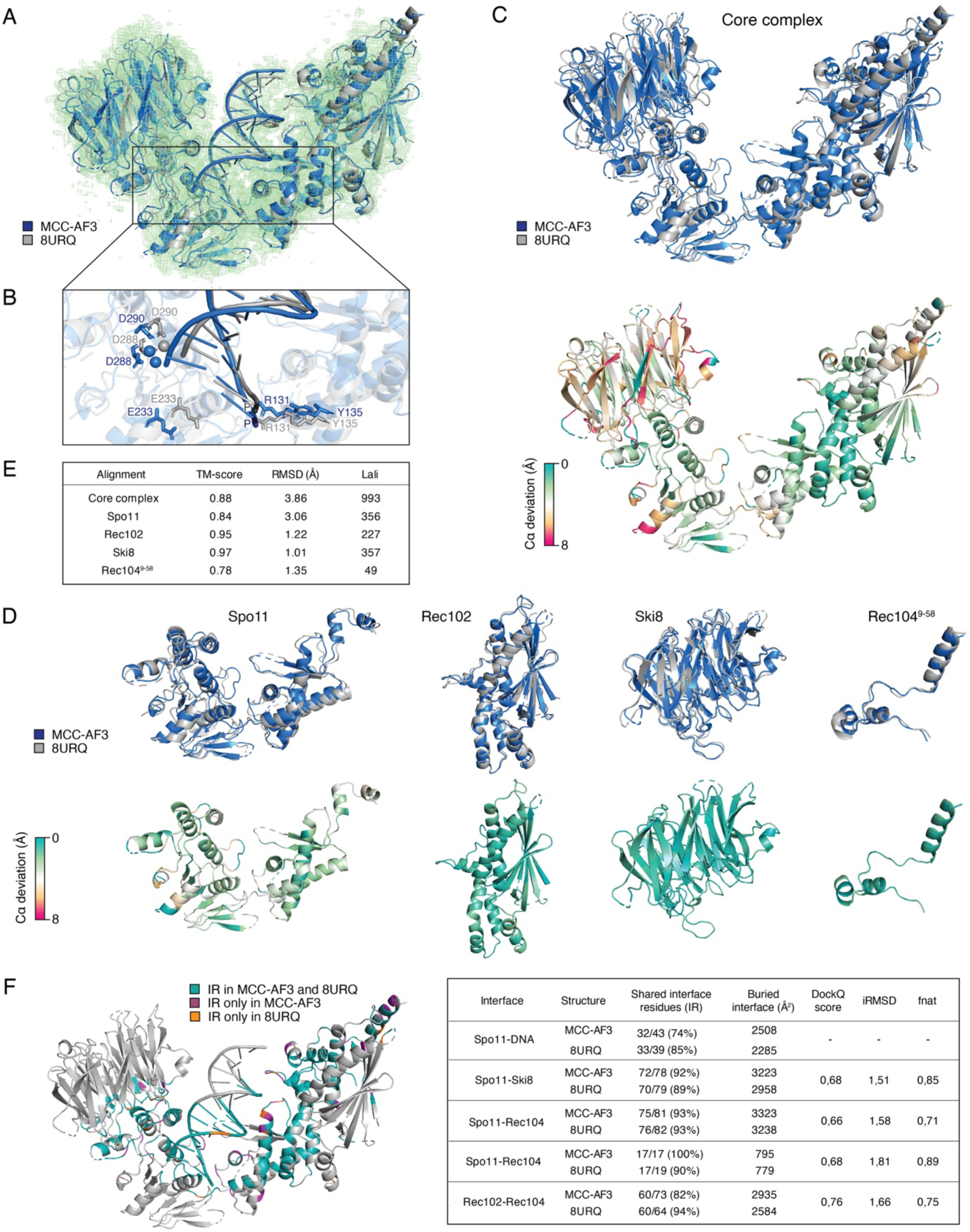
Structural analysis and comparison of the *S. cerevisiae* monomeric core complex predicted by AlphaFold3 (MCC-AF3) and determined by cryo-EM (PDB: 8URQ). (**A-F**) MCC-AF3 corresponds to the Complex 1 of the *S. cerevisiae* dimeric core complex predicted by Alphafold3 (DCC-AF3, Figure 1E) and used throughout this work. MCC-AF3 was trimmed to the resolved regions present in 8URQ. (**A**) Superposition of DNA-bound core complex of MCC-AF3 (blue) and 8URQ (gray) in the electron density map of 8URQ (green, EMDB EMD-42497 nominal 3.3 Å map). (**B**) Active site comparison of MCC-AF3 (blue) and 8URQ (gray). Mg^2+^ ions are shown as spheres. Active site residues (R131, Y135, E233, D288, D290) are shown as sticks. Scissile phosphates (P) are in dark blue (MCC-AF3) or in dark gray (8URQ). (**C**) Top: Superposition of the core complex of MCC-AF3 (blue) and 8URQ (gray). Bottom: residue-wise Cα deviations (0–8 Å) between MCC-AF3 and 8URQ were mapped onto MCC-AF3 for clear visualization. (**D**) Top: per chain superposition of Spo11, Rec102, Ski8, Rec104 from MCC-AF3 (blue) and 8URQ (gray). Bottom: residue-wise Cα deviations (0–8 Å) between MCC-AF3 and 8URQ were mapped onto CC-AF3 for clear visualization. (**E**) Core complex and per chain (Spo11, Rec102, Ski8, Rec104) US-align analysis of MCC-AF3 and 8URQ. MCC-AF3 vs 8URQ shows strong global agreement (TM-score = 0.88) with RMSD = 3.6 Å over 993 aligned residues and per chain agreement with TM-scores 0.78-0.97 and RMSD 1.1-3.1. (**F**) Interface comparison of MCC-AF3 and 8URQ subunits. Interface preservation: MCC-AF3 shares 88% of its total interface residues with 8URQ. On average, the amount of buried interface of MCC-AF3 and 8URQ subunits is similar at 93%. Interfaces were also analyzed using DockQ v2 (Mirabello and Wallner 2024). DockQ analysis over four native interfaces (Spo11-Rec102, Spo11–Ski8, Spo11-Rec104, Rec102-Rec104) yields scores of 0.66–0.75 (overall 0.71), with iRMSD 1.5–1.8 Å, and fnat 0.70–0.89 demonstrating that MCC-AF3 recapitulates the interface geometry observed in 8URQ. TM-score, Template Modeling score; RMSD, root-mean-square deviation; Lali, alignment length; iRMSD, interface RMSD; fnat, fraction of native contacts.

**Supplementary Figure 3:**
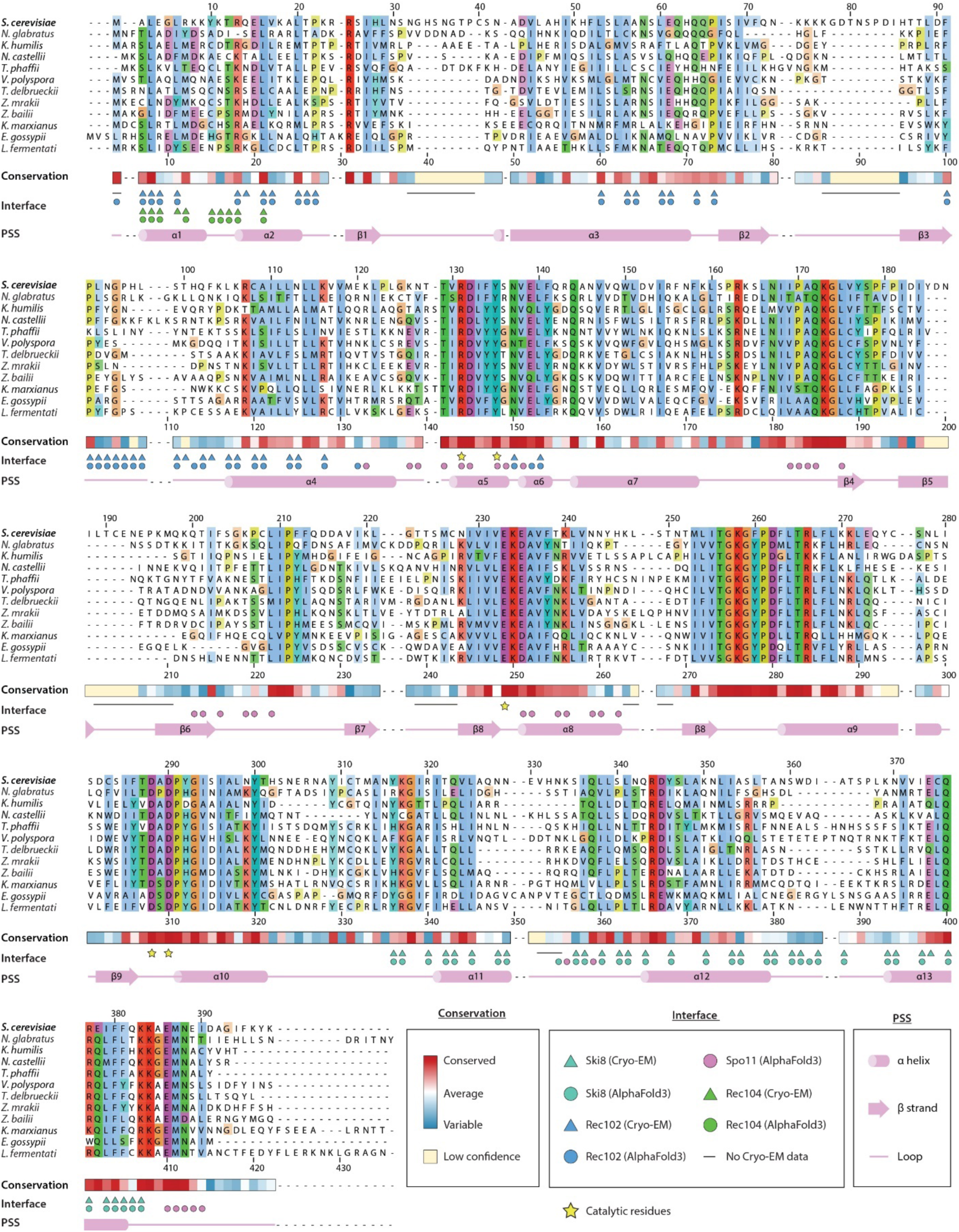
Structure-informed alignment and conservation of Spo11 orthologs. Residues of the Spo11 dimer interface (Figure 2) are indicated by pink circle. UniProt references: *S. cerevisiae* (P23179), *N. glabratus* (Q6FWU6), *K. humilis* (A0AAV5RW59), *N. castellii* (G0VGL2), *T. phaffii* (G8BWK5), *V. polyspora* (A7TTH3), *T. delbrueckii* (G9A007), *Z. mrakii* (A0A7H9B6N5), *Z. bailii* (A0A8J2TC47), *K. marxianus* (W0TCD2), *E. gossypii* (Q75A93), *L. fermentati* (A0A1G4M6H9). The numbering of Spo11 residues from *S. cerevisiae* is above the alignment, and the numbering of aligned residues is below the alignment. Conservation and interface data were calculated as described in Methods. PSS, protein secondary structure.

**Supplementary Figure 4:**
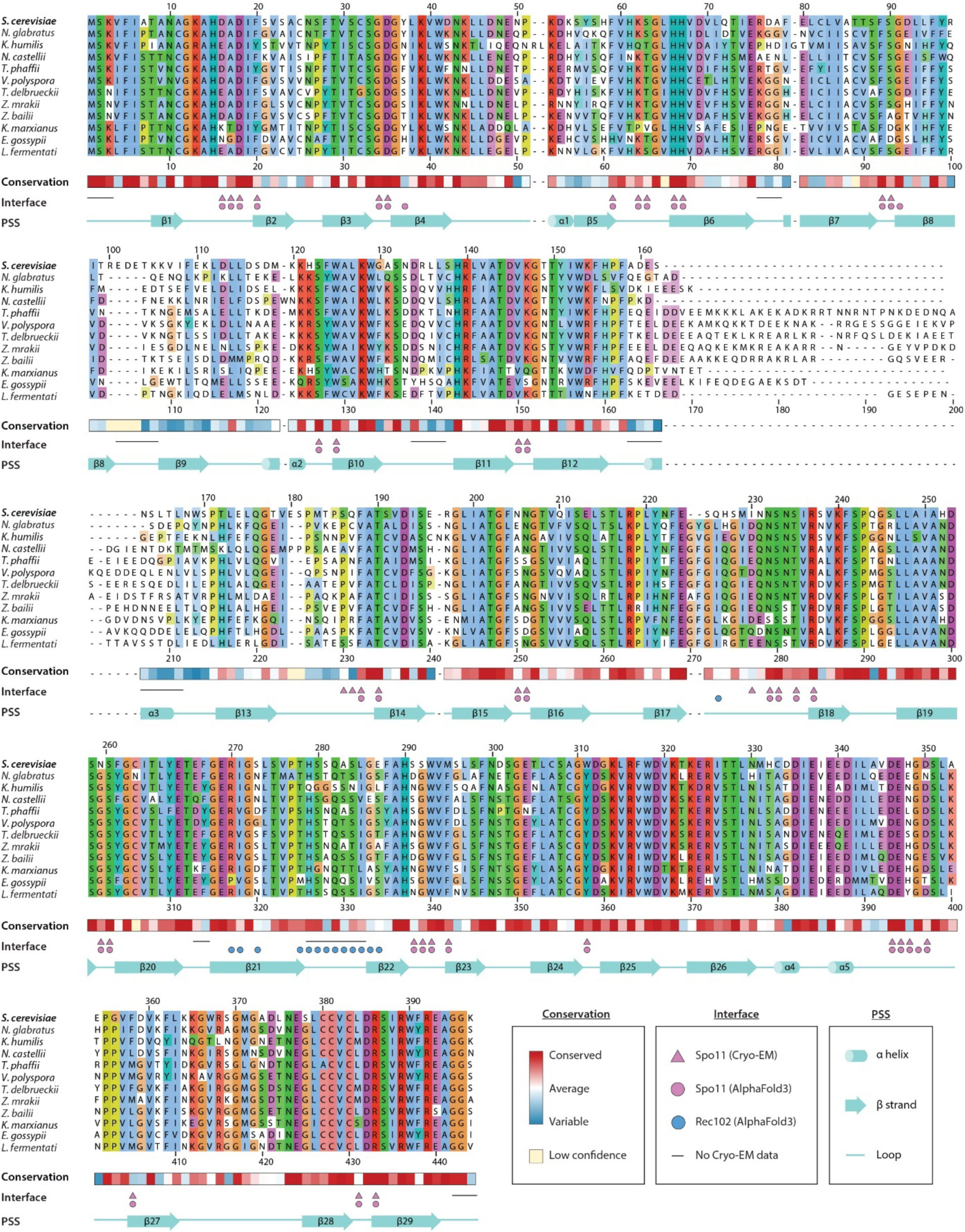
Structure-informed alignment and conservation of Ski8 orthologs. Residues of Ski8 in *trans* contact with Rec102 (Figure 9) are indicated by blue circles. UniProt references: *S. cerevisiae* (Q02793), *N. glabratus* (Q6FQD7), *K. humilis* (A0AAV5RRP0), *N. castellii* (G0VF58), *T. phaffii* (G8BP31), *V. polyspora* (A7TER9), *T. delbrueckii* (G8ZU84), *Z. mrakii* (A0A7H9AYH9), *Z. bailii* (A0A8J2X0D3), *K. marxianus* (W0TCZ4), *E. gossypii* (Q755S9), *L. fermentati* (A0A1G4MG77). The numbering of Ski8 residues from *S. cerevisiae* is above the alignment, and the numbering of aligned residues is below the alignment. Conservation and interface data were calculated as described in Methods. PSS, protein secondary structure.

**Supplementary Figure 5:**
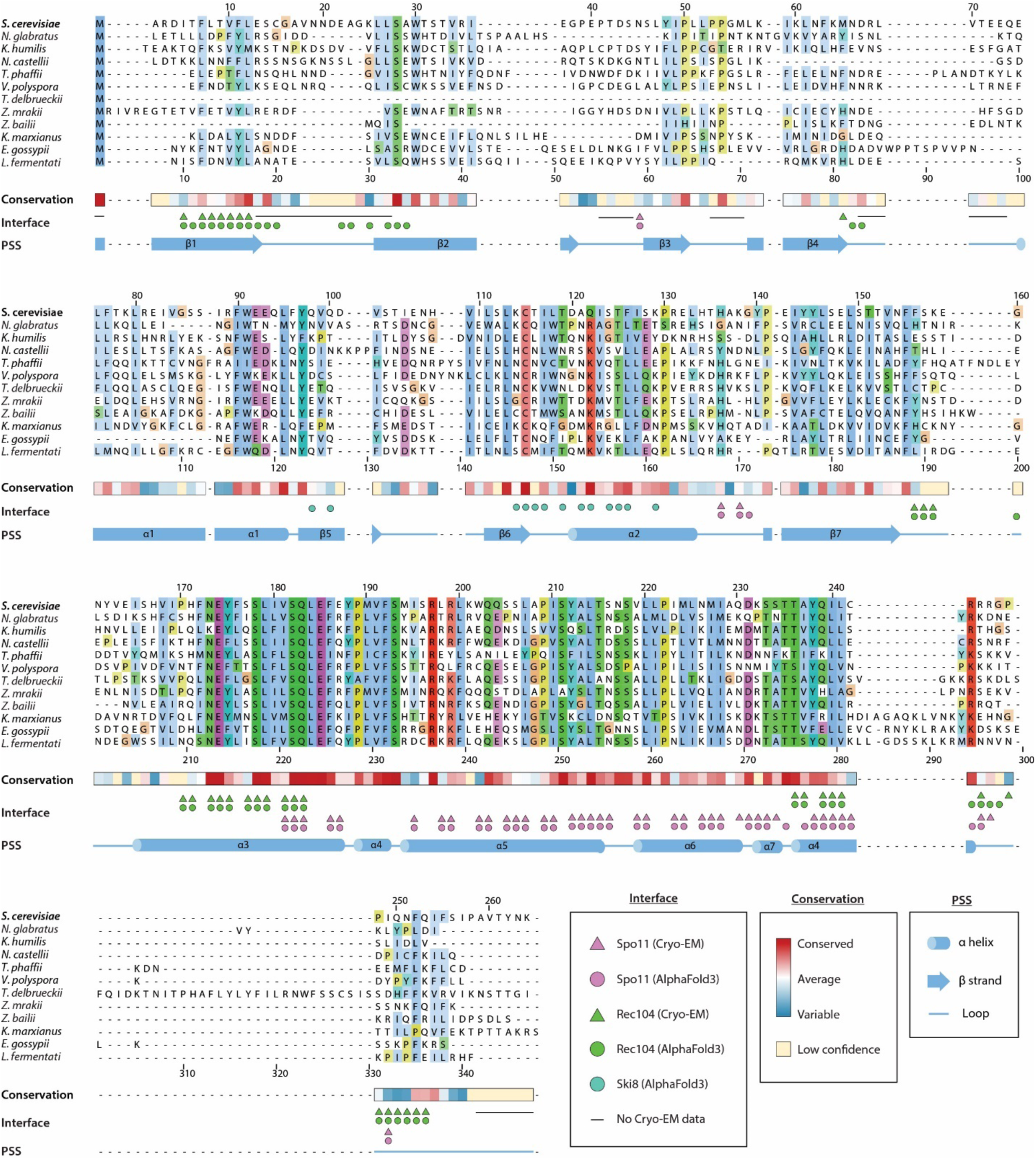
Structure-informed alignment and conservation of Rec102 orthologs. Residues of Rec102 in *trans* contact with Ski8 (Figure 8) are indicated by cyan circles. UniProt references (unless otherwise specified): *S. cerevisiae* (Q02721), *N. glabratus* (A0A0W0C7A9), *K. humilis* (the sequences on Uniprot A0AAV5S006 and NCBI GMM57189.1 are incorrect due to the presence of a mis-annotated intron, which was corrected here), *N. castellii* (G0VKM6), *T. phaffii* (G8BUZ4), *V. polyspora* (A7TKV1), *T. delbrueckii* (G8ZTD8), *Z. mrakii* (A0A7H9B4G4), *Z. bailii* (NCBI: AQZ11626), *K. marxianus* (W0TB81), *E. gossypii* (Q750D8), *L. fermentati* (A0A1G4MAP2). The numbering of Rec102 residues from *S. cerevisiae* is above the alignment, and the numbering of aligned residues is below the alignment. Conservation and interface data were calculated as described in Methods. PSS, protein secondary structure.

**Supplementary Figure 6.**
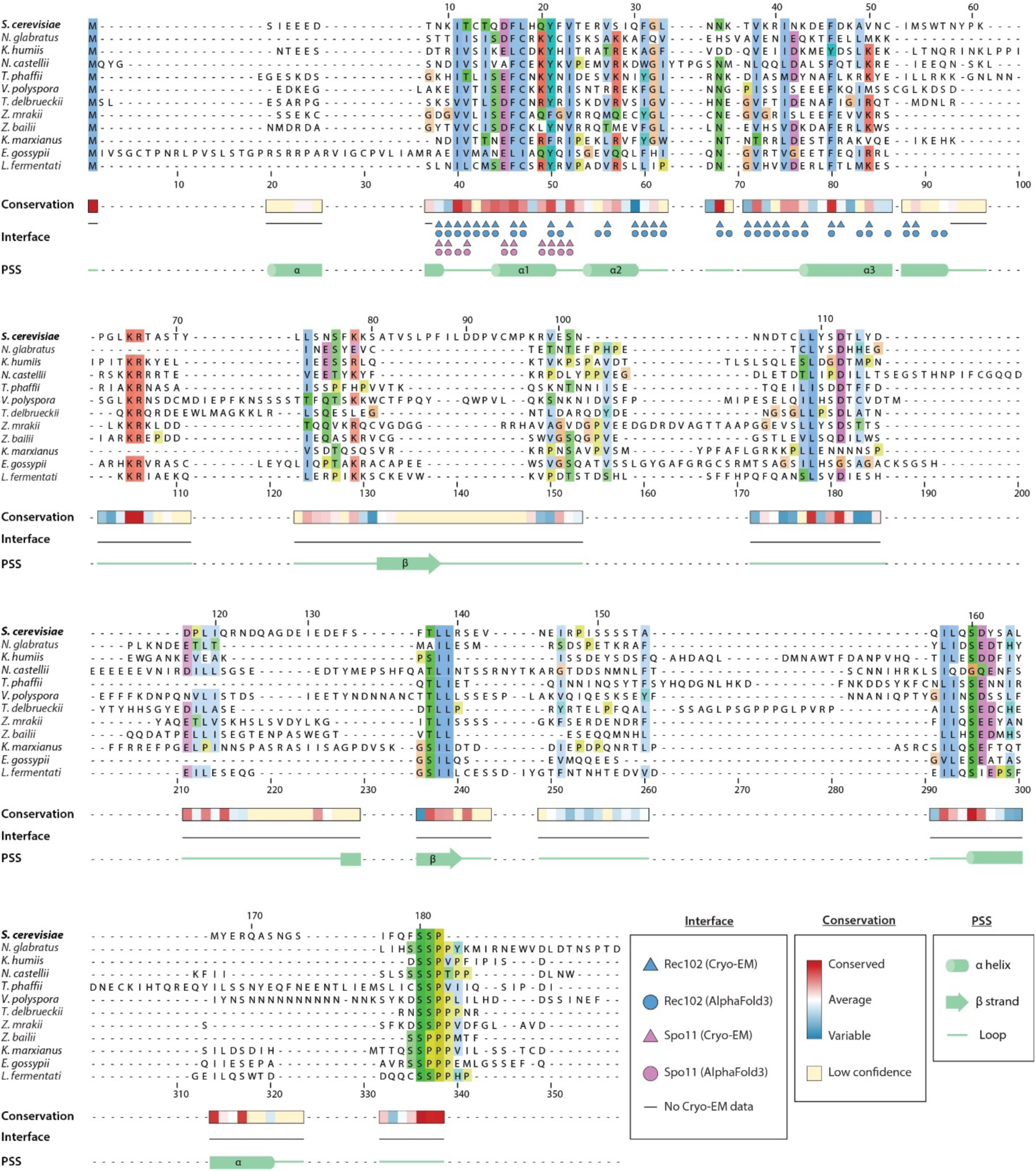
Structure-informed alignment and conservation of Rec104 orthologs. UniProt references (unless otherwise specified): *S. cerevisiae* (P33323), *N. glabratus* (Q6FR03), *K. humilis* (A0AAV5S5B3), *N. castellii* (G0VDU1), *T. phaffii* (G8BNW3), *V. polyspora* (A7TEW4), *T. delbrueckii* (G8ZYK5), *Z. mrakii* (A0A7H9B3P3), *Z. bailii* (NCBI: SJM83633.1), *K. marxianus* (NCBI: BAP72788.1), *E. gossypii* (Q75AR8), *L. fermentati* (A0A1G4MAB8). The numbering of Rec104 residues from *S. cerevisiae* is above the alignment, and the numbering of aligned residues is below the alignment. Conservation and interface data were calculated as described in Methods. PSS, protein secondary structure.

**Supplementary Figure 7:**
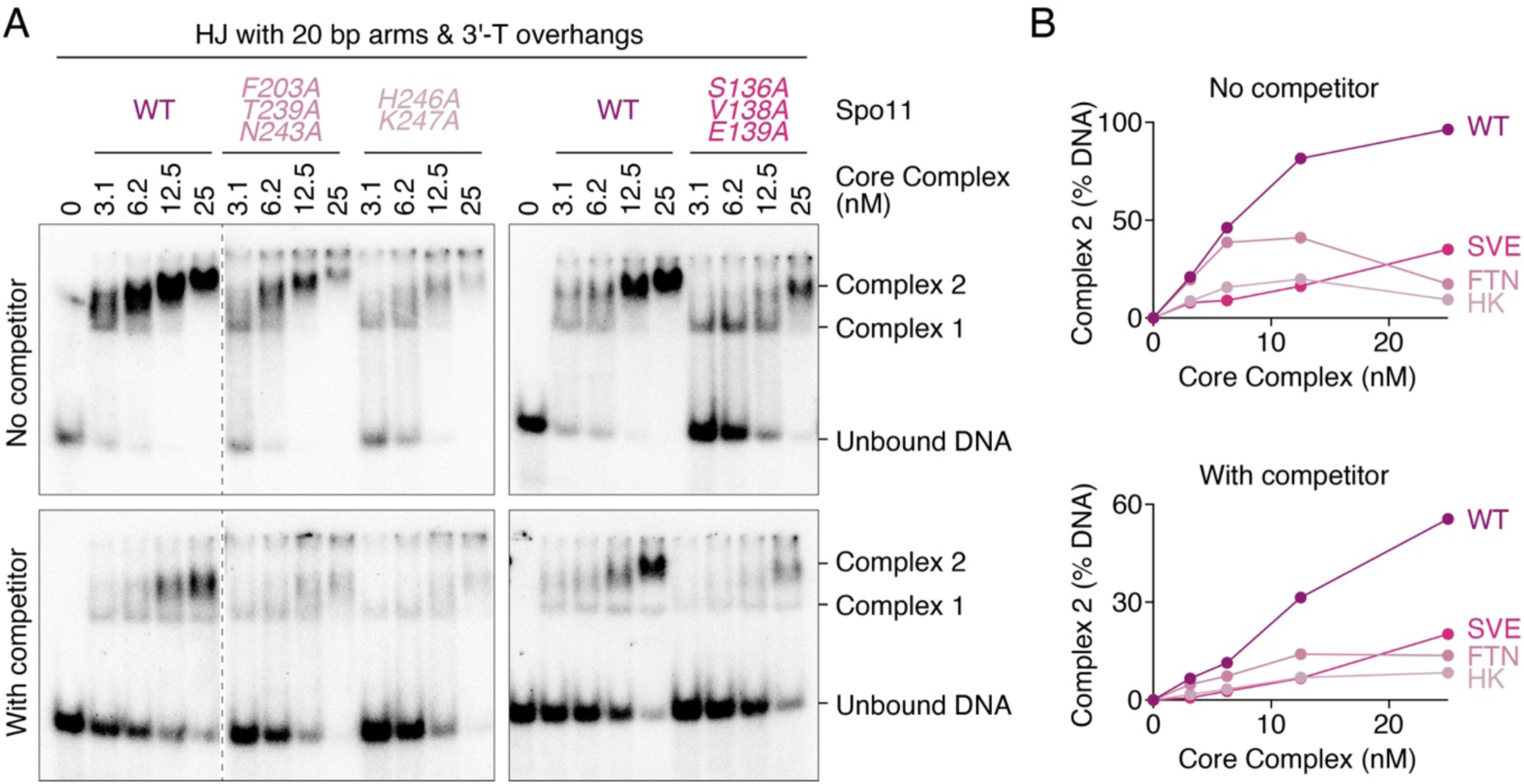
DNA-binding properties of Spo11 dimerization mutants. **(A)** Gel shift analysis of DNA binding by wild-type and mutant core complexes on a radiolabeled Holliday Junction (HJ) substrate (0.5 nM) with or without 20 bp dsDNA competitor (25 nM). **(B)** Quantification of Complex 2 from the gels in panel A. SVE, Spo11-S136A/V138A/E139A; FTN, Spo11-F203A/T239A/N243A; HK, Spo11-H246A/K247A.

**Supplementary Figure 8:**
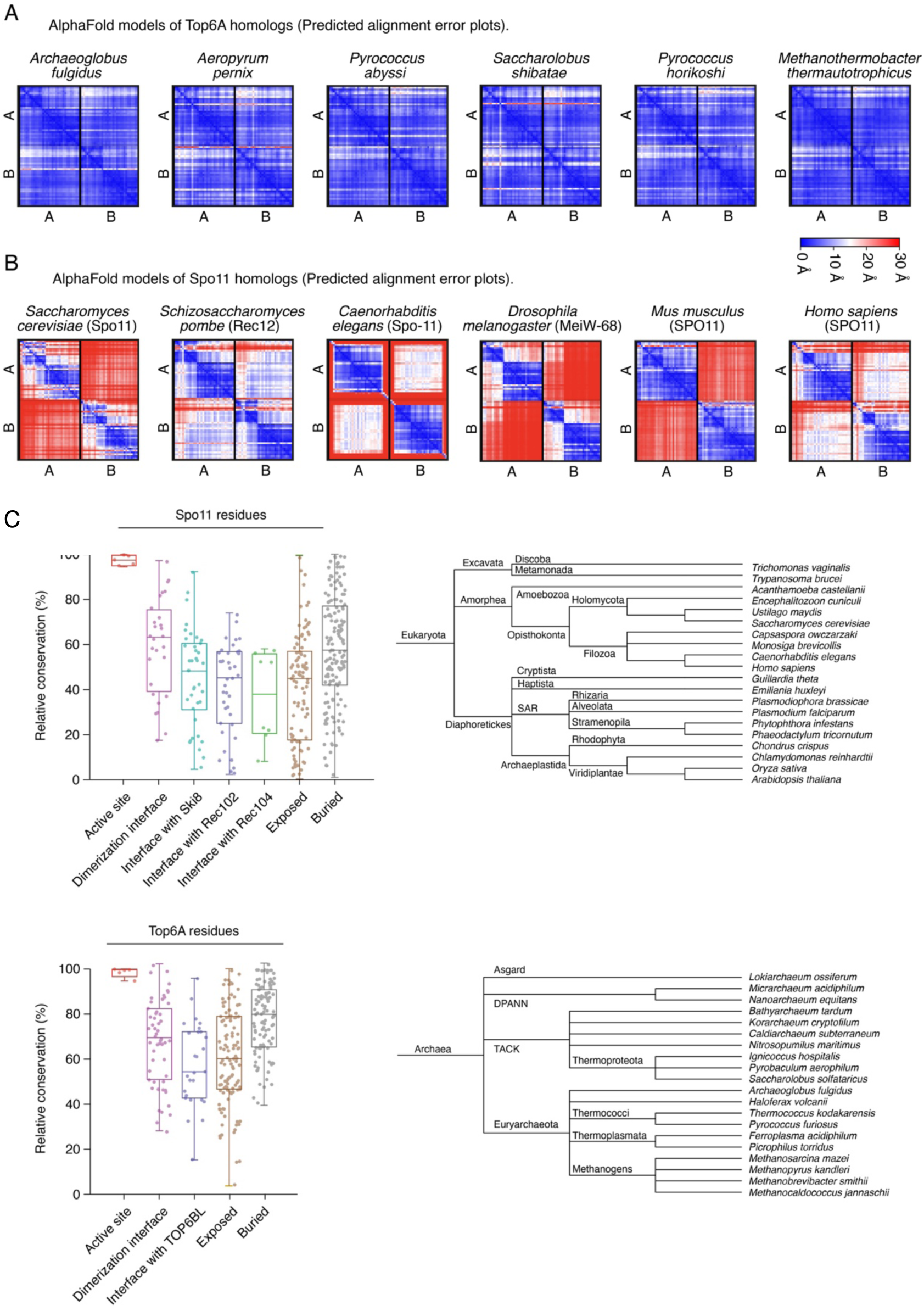
Modeling of Spo11 and Top6A dimers and conservation of dimer interfaces. **(A, B)** Predicted alignment error plots from AlphaFold2 models of Top6A and Spo11 dimers from selected species. The structure and relative position of two Top6A subunits are consistently predicted with high confidence. In contrast, the relative position of two Spo11 subunits (top right and bottom left squares) tend to be predicted with low confidence. **(C)** Sequence conservation analysis of Spo11 proteins across eukaryotes and Top6A proteins across archaea. (Right) Phylogenetic trees of the queried species. (Left) Sequences were aligned to *S. cerevisiae* Spo11 or *M. mazei* Top6A and residues were categorized as exposed, as buried, as parts of the active site or as parts of interaction surfaces based on the AlphaFold model of the core complex (Figure 1E) or a crystal structure of Topo VI (2Q2E, Corbett et al. 2007), respectively.

**Supplementary Figure 9:**
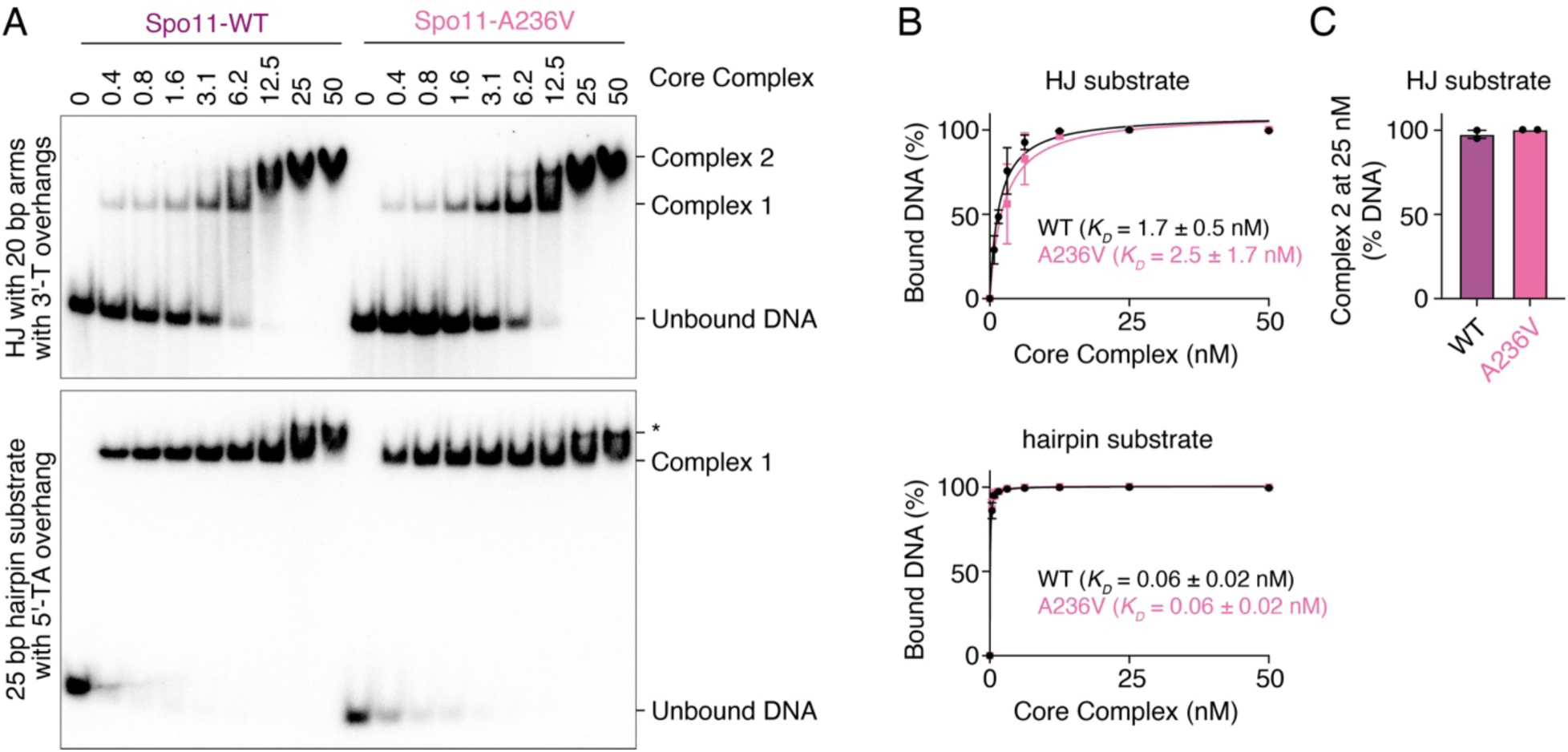
The Spo11-A236V mutant does not affect DNA binding and dimerization. **(A)** Gel shift analysis of the binding activity of wild-type core complexes and Spo11-A236V mutants to Holliday junctions (HJ with 20 bp arms and 1-nt 3′-overhangs), or DNA ends (25 bp hairpin substrates with a 2-nt 5′-overhang). **(B)** Quantification of the affinity of wild-type and A236V mutants for each substrate. **(C)** Quantification of Complex 2 at 25 nM core complexes (mean ± range of two independent experiments).

**Supplementary Figure 10:**
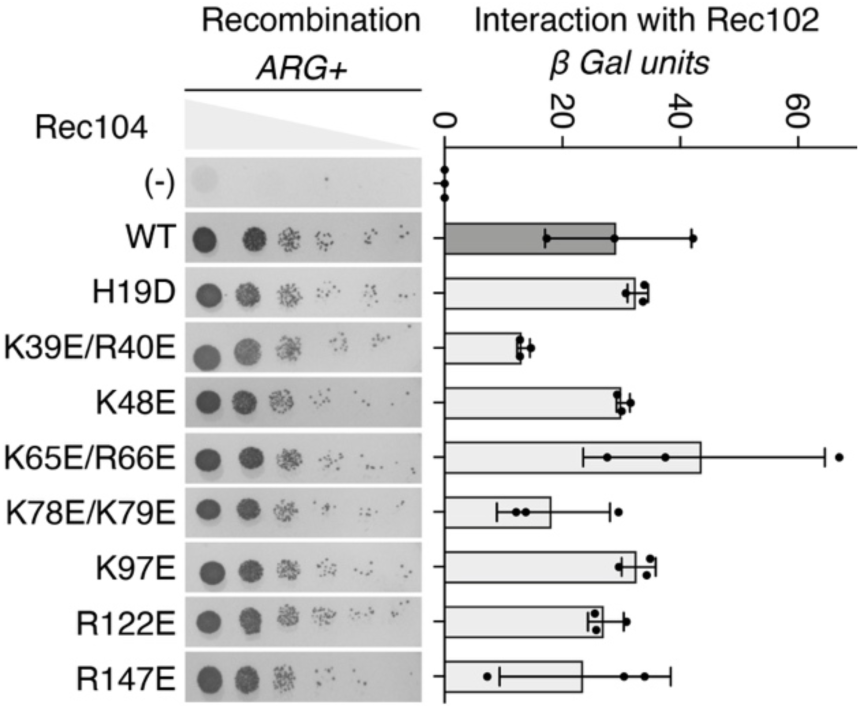
Mutagenesis analysis of positively charged Rec104 residues. Effects of Rec104 mutations on (left) recombination established by *arg4* heteroallele recombination assay and on (right) the interaction with Rec102 established by yeast two-hybrid assay (mean ± SD of three biological replicates).

**Supplementary Figure 11:**
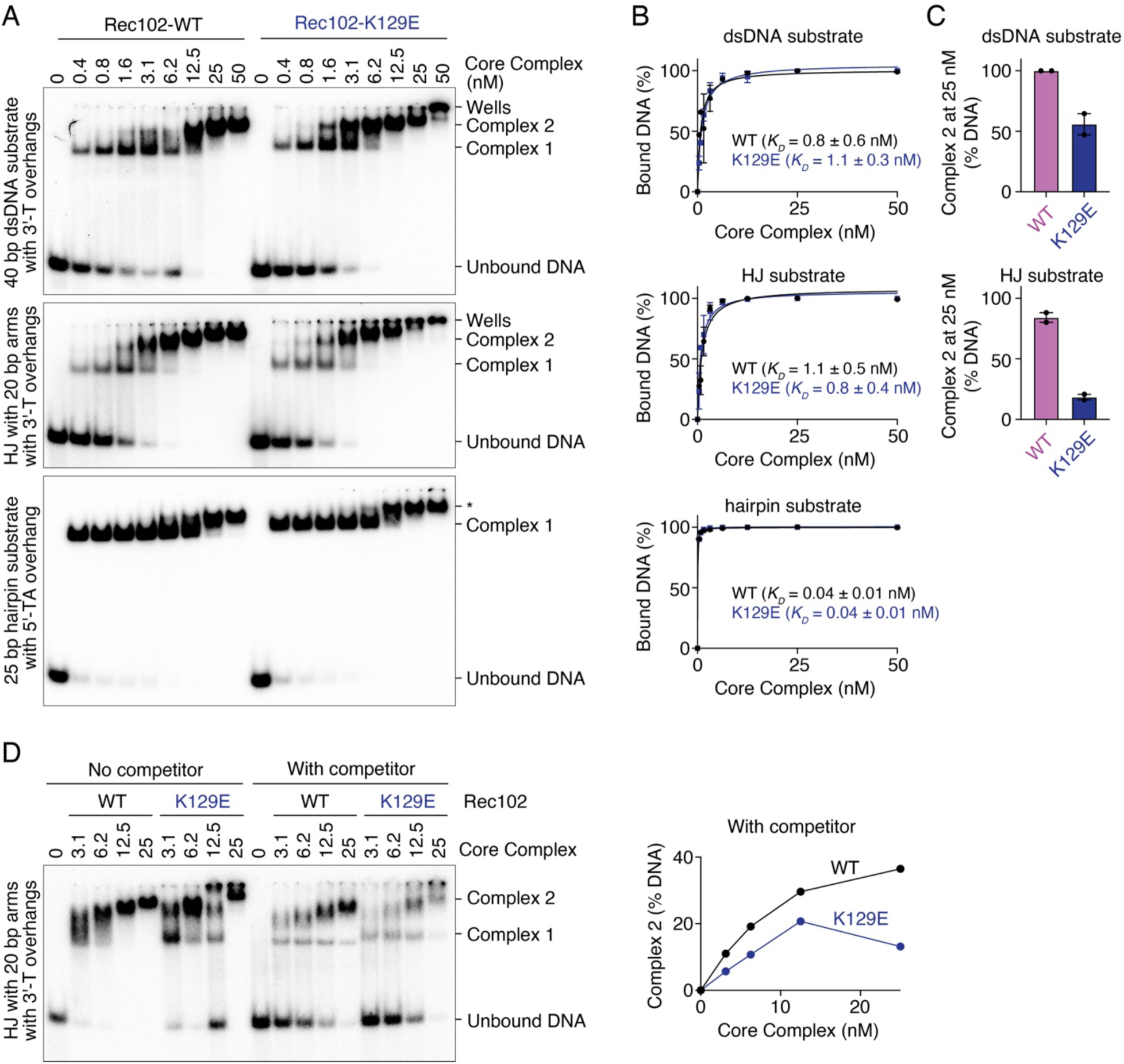
Impact of Rec102-K129E mutation on the DNA-binding activity of the core complex. **(A)** Gel shift analysis of the binding activity of wild-type core complexes and Rec102-K129E mutants to DNA duplexes (40 bp dsDNA with 1-nt 3′-overhangs), Holliday junctions (HJ with 20 bp arms and 1-nt 3′-overhangs), or DNA ends (25 bp hairpin substrates with a 2-nt 5′-overhang). **(B)** Quantification of the affinity of wild-type and K129E mutants for each substrate. **(C)** Quantification of Complex 2 at 25 nM core complexes (mean ± range of two independent experiments). **(D)** Gel shift analysis of binding to a labeled HJ substrate (0.5 nM) in the presence or absence of a 20 bp dsDNA competitor (25 nM). The assembly of Complex 2 in the presence of competitor is plotted.

**Supplementary Figure 12.**
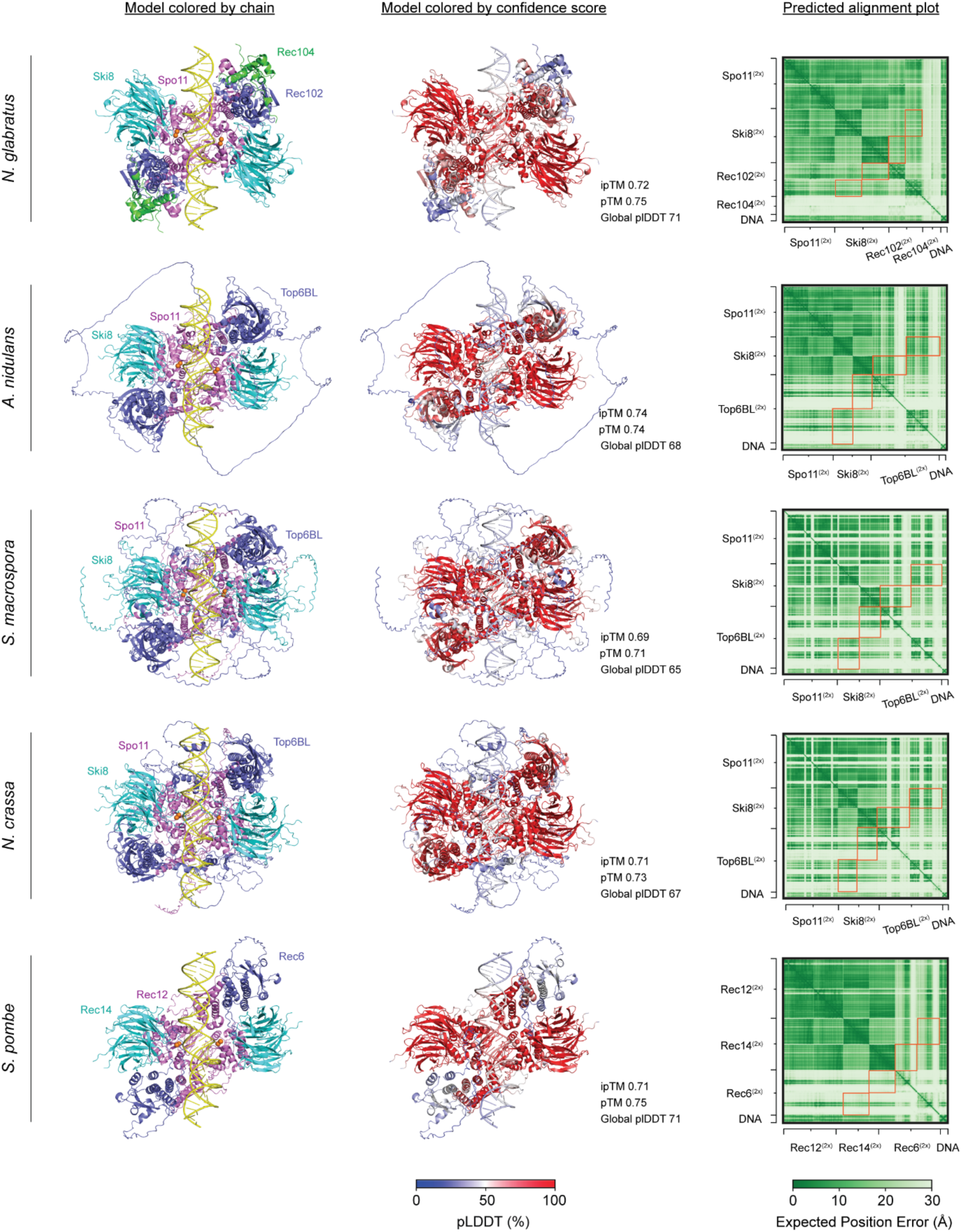
Structural modeling of Spo11 core complexes in different ascomycetes. AlphaFold3 models of DNA-bound dimeric Spo11 core complexes from *Nakaseomyces glabratus* (Q6FWU6, A0A0W0C7A9, Q6FR03, Q6FQD7), *Aspergillus nidulans* (Q5ATX1, Q5BDJ3, Q5B4V3) *Sordaria macrospora* (Q6WRU4, F7VPQ4 re-annotated, Q6URC5), *Neurospora crassa* (Q1K767, Q1K546, Q7RVN5) and *Schizosaccharomyces pombe* (P40384, P40385, Q09150). Left: AlphaFold3 models colored by chain. Spo11 homologs are shown in pink; Ski8 homologs are shown in cyan; Rec102 homologs and the transducer domain of TOP6BL homologs are shown in blue; Rec104 homologs are shown in green, DNA is shown in yellow and Mg^2+^ ions are in orange. Middle: AlphaFold3 models colored by confidence score. Right: predicted alignment error plot. Despite extensive variability in the sequence and structure of Rec102 and Top6BL homologs, all the models show Ski8-Rec102 or Ski8-Top6BL interactions, consistent with our interpretation that this interaction is an important pre-requisite for DNA cleavage in fungi. The confidence in the structure and relative position of Ski8-Rec102 and Ski8-Top6BL interactions is indicated by orange squares on the predicted alignment error plots.

### Supplementary Tables

**Supplementary Table 1:**
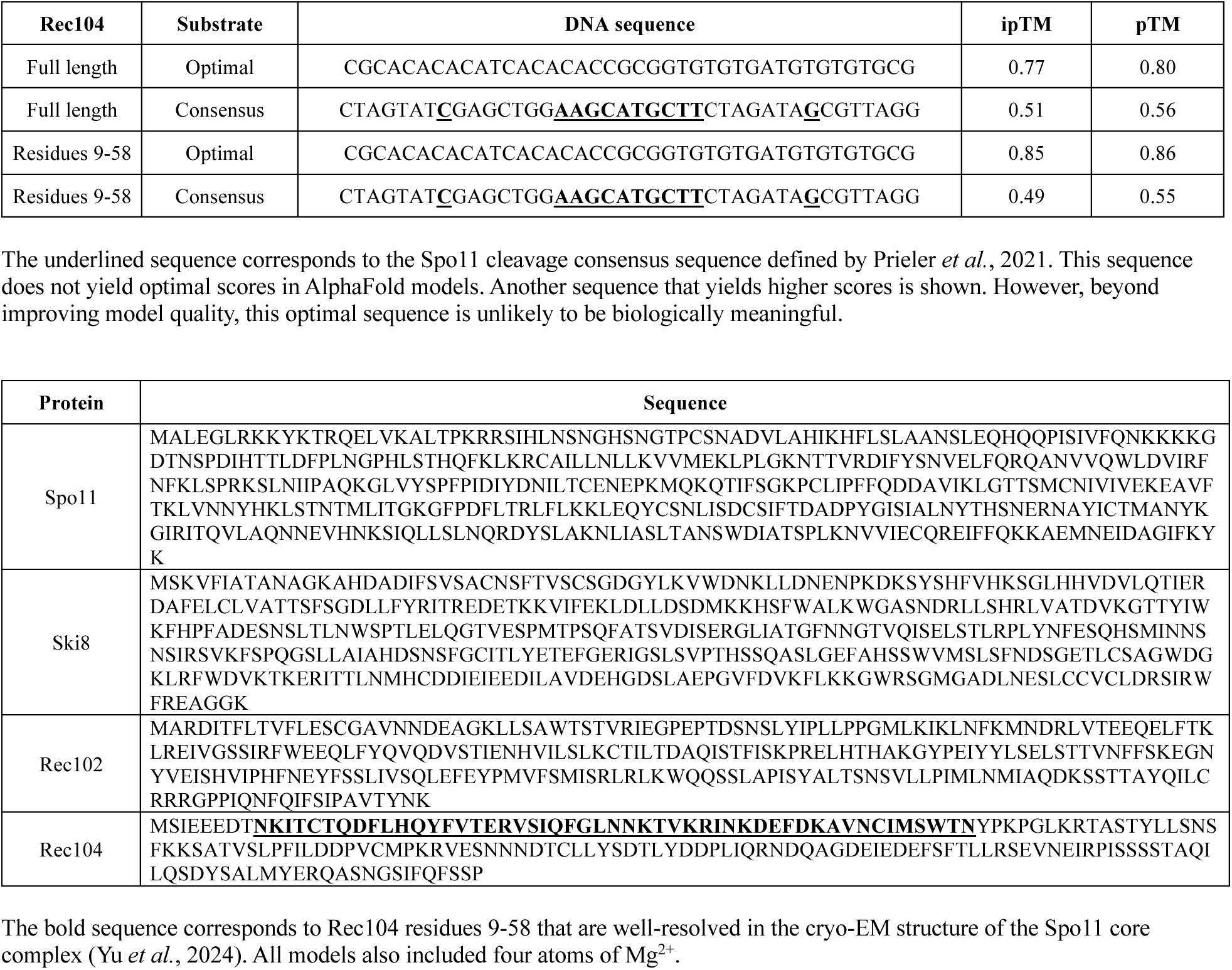
Input sequences and quality scores of AlphaFold models.

**Supplementary Table 2:** List of plasmids.

| Plasmid | Description | Reference |
| --- | --- | --- |
| pCCB587 | Ski8 in pFastBac1 | (Claeys Bouuaert et al. 2021b) |
| pCCB588 | Rec102 in pFastBac1 | (Claeys Bouuaert et al. 2021b) |
| pCCB589 | Rec104 in pFastBac1 | (Claeys Bouuaert et al. 2021b) |
| pCCB592 | Spo11-HisFlag in pFastBac1 | (Claeys Bouuaert et al. 2021b) |
| pCCB615 | Ski8-HisFlag in pFastBac1 | (Claeys Bouuaert et al. 2021b) |
| pCCB617 | MBP-Rec102 in pFastBac1 | (Claeys Bouuaert et al. 2021b) |
| pCCB758 | REC102::NatMX6 cassette in Topo Blunt II vector | This study |
| pCCB759 | REC104::NatMX6 in Topo blunt vector | This study |
| pCCB767 | Spo11-HisFlag:HphMX4 cassette in pRS314 | This study |
| pCCB771 | Spo11-T239A/K240A in pCCB767 | This study |
| pCCB772 | Spo11-K126A/N127A in pCCB767 | This study |
| pHAB001 | MBP-Tev-Rec102 in pFastBac1 | This study |
| pHAB008 | Rec104-H19D in pSK293 (Y2H vector) | This study |
| pHAB009 | Rec104-K39E/R40E in pSK293 (Y2H vector) | This study |
| pHAB010 | Rec104-K48E in pSK293 (Y2H vector) | This study |
| pHAB011 | Rec104-K65E/R66E in pSK293 (Y2H vector) | This study |
| pHAB012 | Rec104-K78E/K79E in pSK293 (Y2H vector) | This study |
| pHAB013 | Rec104-K97E in pSK293 (Y2H vector) | This study |
| pHAB014 | Rec104-R122E in pSK293 (Y2H vector) | This study |
| pHAB015 | Rec104-R147E in pSK293 (Y2H vector) | This study |
| pHAB016 | Rec102-K79E/R81E in pSK282 (Y2H vector) | This study |
| pHAB017 | Rec102-K79E in pSK282 (Y2H vector) | This study |
| pHAB018 | Rec102-R81E in pSK282 (Y2H vector) | This study |
| pHAB019 | Rec102-R89E in pSK282 (Y2H vector) | This study |
| pHAB020 | Rec102-K129E in pSK282 (Y2H vector) | This study |
| pHAB021 | Rec102-H134A in pSK282 (Y2H vector) | This study |
| pHAB022 | Rec102-H136A in pSK282 (Y2H vector) | This study |
| pHAB023 | Rec102-K138E in pSK282 (Y2H vector) | This study |
| pHAB024 | Rec102-K158E in pSK282 (Y2H vector) | This study |
| pHAB025 | Rec102-H171D in pSK282 (Y2H vector) | This study |
| pHAB026 | Rec102-R197E in pSK282 (Y2H vector) | This study |
| pHAB027 | Rec102-R199E/K201E in pSK282 (Y2H vector) | This study |
| pHAB028 | Rec102-K232E in pSK282 (Y2H vector) | This study |
| pHAB029 | Rec102-R243E/R244E/R245E in pSK282 (Y2H vector) | This study |
| pHAB030 | Rec102-R68E in pSK282 (Y2H vector) | This study |
| pHAB031 | Rec102-K64E in pSK282 (Y2H vector) | This study |
| pHAB032 | Rec102-K60E in pSK282 (Y2H vector) | This study |
| pHAB033 | Rec102-K58E in pSK282 (Y2H vector) | This study |
| pHAB054 | MBP-tev-mVenus-Rec102 in pFastBac1 | This study |
| pHAB062 | Rec102-K129E in pCCB758 | This study |
| pHAB066 | Rec102-R243E/R244E/R245E in pCCB758 | This study |
| pHAB086 | Rec102-R245E in pCCB758 | This study |
| pHAB087 | MBP-tev-Rec102-R245E in pFastBac1 | This study |
| pHAB092 | Spo11-S136A/V138A/E139A in pCCB767 | This study |
| pHAB093 | Spo11-F203A in pCCB767 | This study |
| pHAB094 | Spo11-F203A/K206A/P207A in pCCB767 | This study |
| pHAB095 | Spo11-T239A/N243A in pCCB767 | This study |
| pHAB103 | MBP-tev-Rec102-R243E/R244E/R245E in pFastBac1 | This study |
| pHAB119 | Spo11-A236V-HisFlag in pFastBac1 | This study |
| pHAB124 | Spo11-S136A/V138A/E139A-HisFlag in pFastBac1 | This study |
| pHAB125 | Spo11-F203A-HisFlag in pFastBac1 | This study |
| pHAB156 | MBP-tev-Rec102-K129E in pFastBac1 | This study |
| pHAB160 | Spo11-F203A/T239A/N243A-HisFlag in pFastBac1 | This study |
| pHAB161 | Spo11-F203A/T239A/N243A in pCCB767 | This study |
| pHAB162 | Spo11-S136A/V138A/E139A/F203A in pCCB767 | This study |
| pHAB164 | Spo11-F203A/K206A in pCCB767 | This study |
| pHAB170 | Spo11-K240A in pCCB767 | This study |
| pHAB171 | Spo11-H246A/K247A in pCCB767 | This study |
| pHAB177 | Spo11-H246A/K247A-HisFlag in pFastBac1 | This study |
| pHAB202 | Rec102-R243E in pCCB758 | This study |
| pHAB203 | Rec102-R244E in pCCB758 | This study |
| pRS314 | Yeast shuttle vector (*CEN6 ARS4 TRP1*) | (Sikorski and Hieter 1989) |
| pSK272 | LexA empty Y2H vector | (Arora et al. 2004) |
| pSK276 | Gal4AD empty Y2H vector | (Arora et al. 2004) |
| pSK282 | LexA-Rec102 Y2H vector (pCA1-Rec102) | (Arora et al. 2004) |
| pSK293 | Rec104-LexA Y2H vector (pCA11-Rec104) | (Maleki et al. 2007) |
| pSK302 | Gal4AD-Rec102 Y2H vector (pACT2-Rec102) | (Maleki et al. 2007) |
| pSK310 | Rec104-Gal4AD Y2H vector (pATC2-2-Rec104) | (Arora et al. 2004) |
| pSK806 | SPO11-HisFlag::HphMX6 cassette in pUC19 | (Claeys Bouuaert et al. 2021b) |

**Supplementary Table 3:**
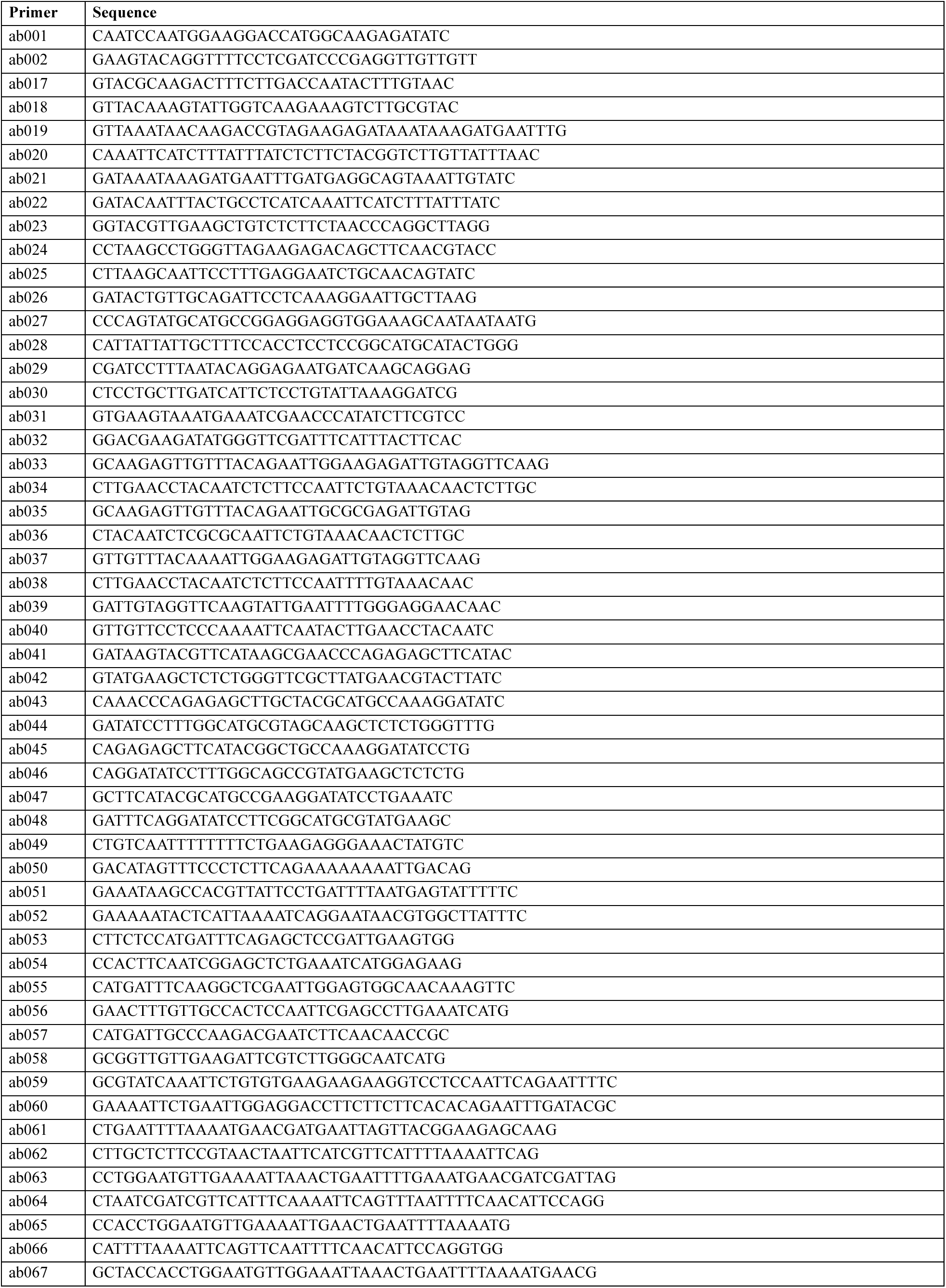
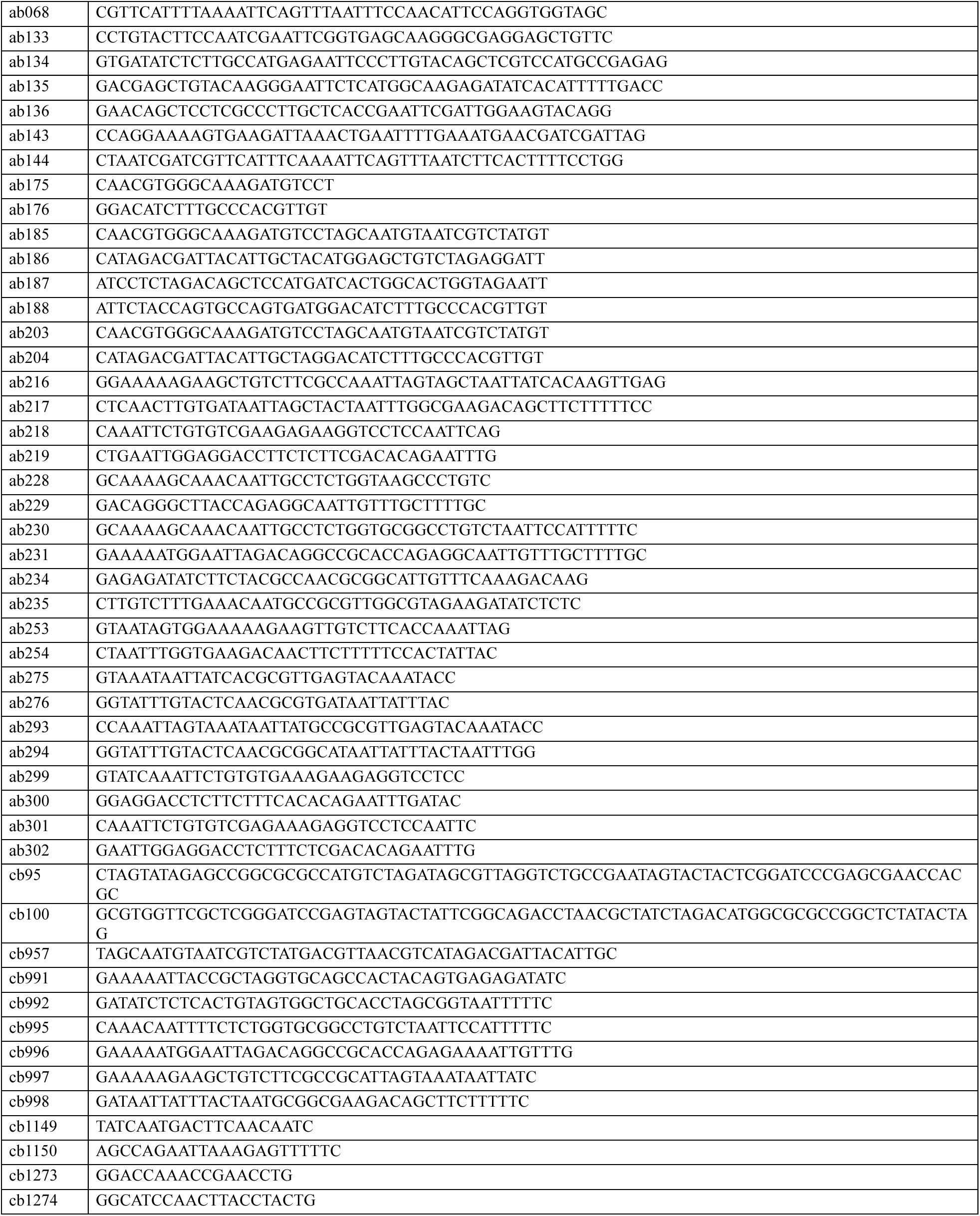
List of oligonucleotides.

**Supplementary Table 4:**
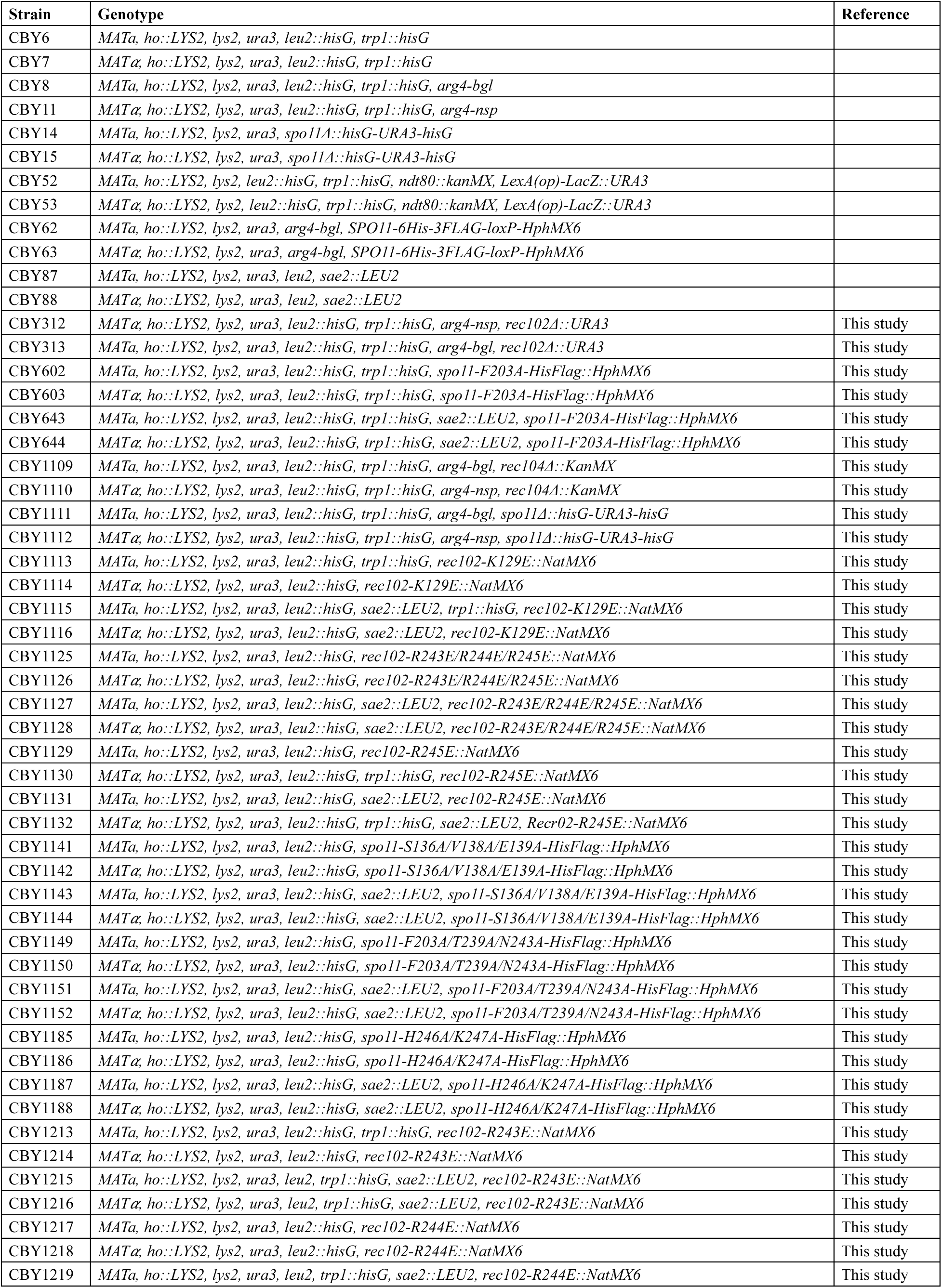
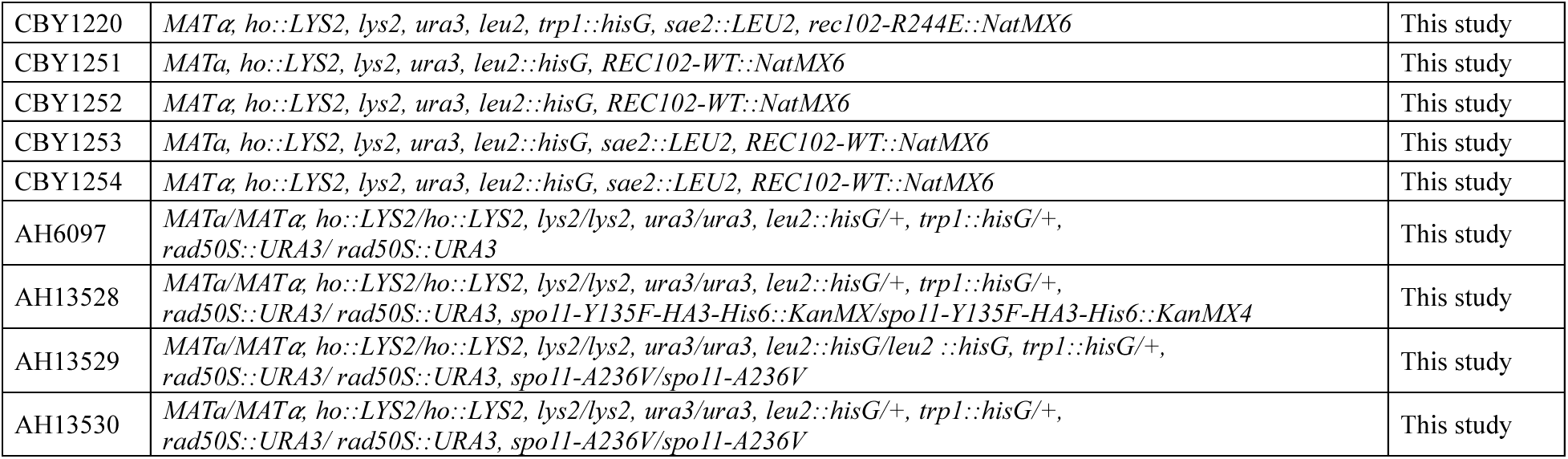
List of strains.

## Notes

### Competing Interest Statement

The authors have declared no competing interest.

